# Energy landscapes reveal agonist control of GPCR activation via microswitches

**DOI:** 10.1101/627026

**Authors:** Oliver Fleetwood, Pierre Matricon, Jens Carlsson, Lucie Delemotte

## Abstract

Agonist binding to G protein-coupled receptors (GPCRs) leads to conformational changes in the transmembrane region that activate cytosolic signaling pathways. Although high resolution structures of different receptor states are available, atomistic details of the allosteric signalling across the membrane remain elusive. We calculated free energy landscapes of the *β*_2_ adrenergic receptors activation using atomistic molecular dynamics simulations in an optimized string of swarms framework, which sheds new light on how microswitches govern the equilibrium between conformational states. Contraction of the extracellular binding site in the presence of the agonist BI-167107 is obligatorily coupled to conformational changes in a connector motif located in the core of the transmembrane region. The connector is probabilistically coupled to the conformation of the intracellular region. An active connector promotes desolvation of a buried cavity, a twist of the conserved NPxxY motif, and an interaction between two conserved tyrosines in transmembrane helices 5 and 7 (Y-Y motif), which leads to a larger population of active-like states at the G protein binding site. This coupling is augmented by protonation of the strongly conserved Asp79^2.50^. The agonist binding site hence communicates with the intracellular region via a cascade of locally connected microswitches. Characterization of these can be used to understand how ligands stabilize distinct receptor states and contribute to development drugs with specific signaling properties. The developed simulation protocol is likely transferable to other class A GPCRs.

**Graphical TOC Entry:** 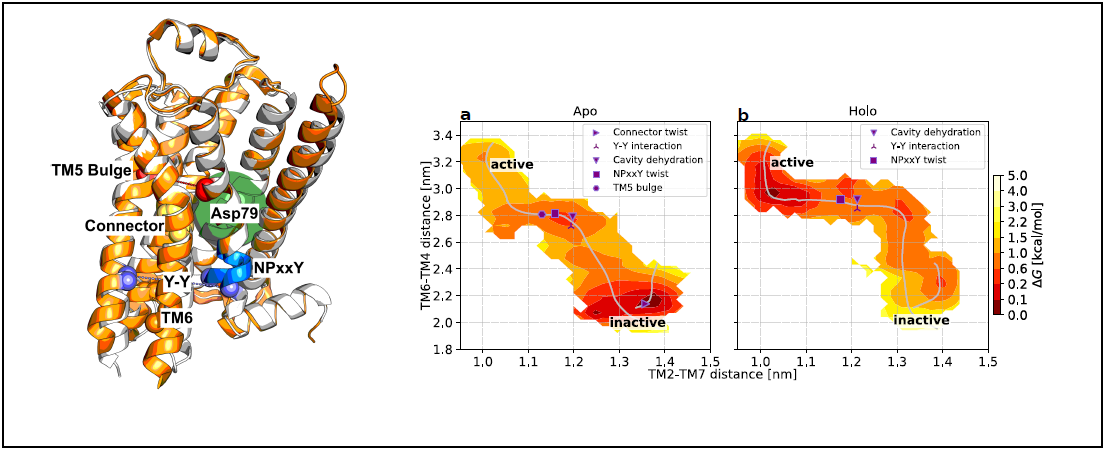

## Introduction

G protein-coupled receptors (GPCRs) are membrane proteins that activate cellular signaling pathways in response to extracellular stimuli. There are more than 800 GPCRs in the human genome (*1*) and these recognize a remarkably large repertoire of ligands such as neurotransmitters, peptides, proteins, and lipids. This large superfamily plays essential roles in numerous physiological processes and has become the most important class of drug targets (*2*). All GPCRs share a common architecture of seven transmembrane (TM) helices, which recognizes the cognate ligand in the extracellular region and triggers intracellular signals via a more conserved cytosolic domain (Fig. 1) (*3, 4*). GPCRs are inherently flexible proteins that exist in multiple conformational states and drug binding alters the relative populations of these. Agonists will shift the equilibrium towards active-like receptor conformations, which promote binding of G proteins and other cytosolic proteins (e.g. arrestin), leading to initiation of signaling via multiple pathways. In the apo state, GPCRs can still access active-like conformations and thereby exhibit a smaller degree of signaling, which is referred to as basal activity.

**Figure 1:**
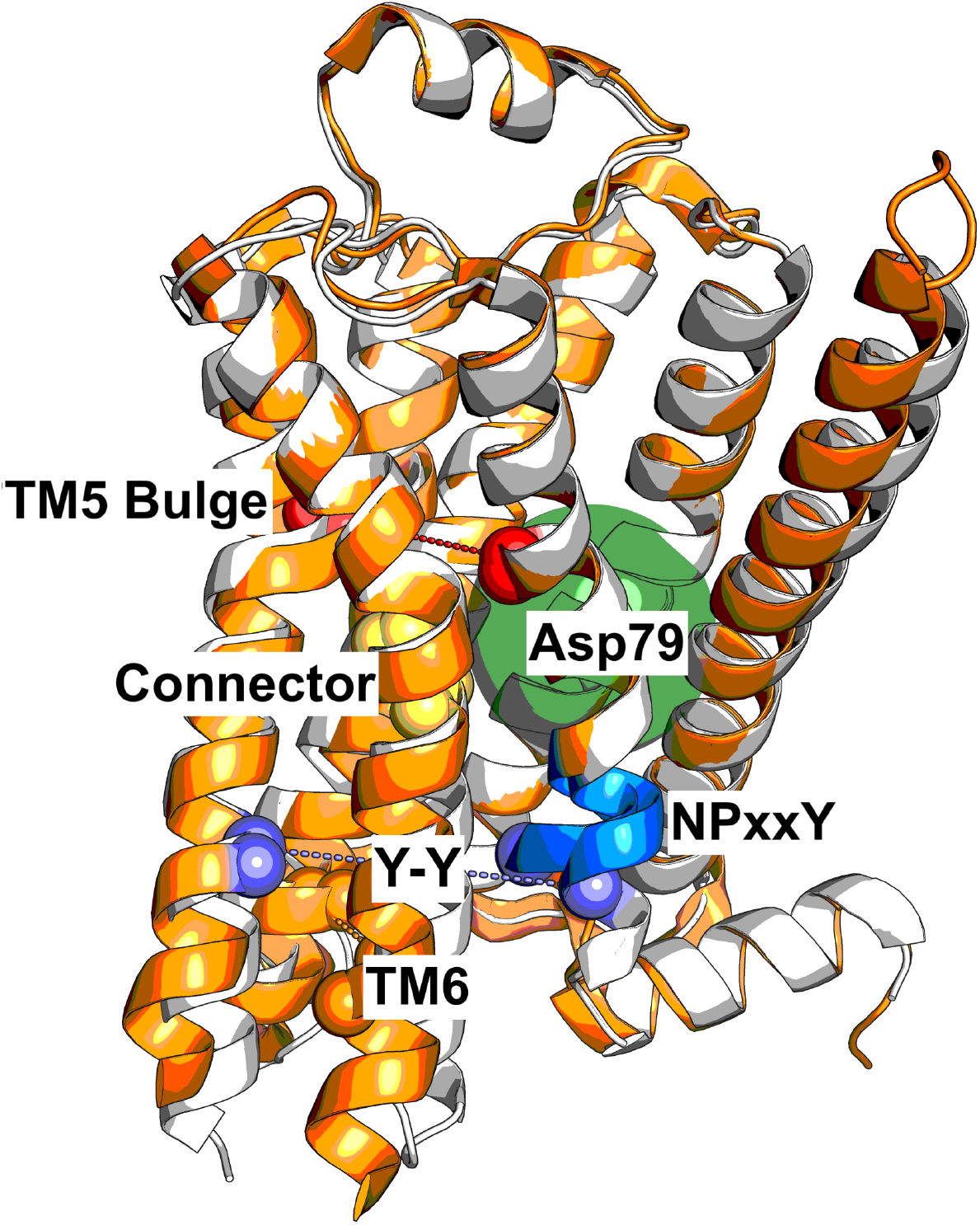
The activation mechanism of GPCRs involves a series of microswitches. Binding of the agonist BI-167107 leads to an inward bulge of TM5 (quantified as the distance between Ser207^5.46^ and Gly315^7.41^, red spheres), which leads to a conformational change in the connector region (Ile121^3.40^ and Phe282^6.44^, yellow spheres). The transmembrane cavity surrounding Asp79^2.50^ dehydrates (filled green circle) and the NPxxY motif (blue cartoon) twists upon activation, leading the Y-Y interaction (Tyr326^7.53^ and Tyr219^5.58^, purple spheres). TM6 moves outwards to create the G protein binding site. The inactive (PDB ID 2RH1 (*5*)) and active (PDB ID 3P0G (*6*)) structures are represented in white and orange, respectively.

Breakthroughs in structure determination of GPCRs during the last decade have provided insights into the process of activation at atomic resolution (Fig. 1). In particular, crystal structures of the *β*_2_ adrenergic receptor (*β*_2_AR) in both active and inactive conformational states (*5–14*) have revealed hallmarks of GPCR activation. The observations made for this prototypical receptor have recently been reinforced by crystal and cryogenic electron microscopy (cryo-EM) structures for other family members (*15*). The most prominent features of GPCR activation are a large ∼ 1.0 − 1.4 nm outward movement of TM6 and a slight inward shift of TM7 on the intracellular side (Fig. 1), which create a large cavity for binding of cytosolic proteins.

Conserved changes in the extracellular part are more difficult to discern due to the lower sequence conservation in this region. In general, the structural changes are relatively subtle and only involve a small contraction of the orthosteric site (*5–7*). In the case of the *β*_2_AR, the catechol group of adrenaline forms hydrogen bonds with Ser207^5.46^ (superscripts denote Ballesteros-Weinstein numbering (*16*)), which leads to a ∼ 0.2 nm inward bulge of TM5. These structural changes then propagate through the receptor via several conserved motifs. The rearrangement of TM5 influences a connector region (PI^3.40^F^6.44^ motif), which is in contact with the highly conserved Asp79^2.50^ and NP^7.50^xxY^7.53^ motif via a network of ordered water molecules. A water-filled cavity surrounding Asp79^2.50^ contributes to stabilizing a sodium ion in several crystal structures of the inactive state, e.g. for the *β*_1_AR (*17, 18*). Upon activation, the cavity collapses, leading to dehydration and displacement of the sodium ion and potentially protonation of Asp79^2.50^ (*19, 20*). Activation also involves a twist of the NPxxY motif, which reorients Tyr326^7.53^ to a position where it can form a water-mediated interaction with Tyr219^5.58^ (Y-Y interaction) (*21*) and together with the displacement of TM6 enable formation of the G protein binding site. Characterization of the role these individual microswitches play in determining the conformational ensemble of structures could guide the design of drugs with tailored signaling profiles.

The allosteric control of GPCR activation by extracellular ligands cannot be fully understood from the static structures captured by crystallography or cryo-EM. Mutagenesis and spectroscopy studies (*22–25*) have suggested that the efficacy of a ligand is determined by a complex interplay between different microswitches and population of distinct states lead to specific functional outcomes. Molecular dynamics (MD) simulations are well suited to study the conformational landscape of GPCRs as this method can provide an atomistic view of the flexible receptor in the presence of membrane, aqueous solvent, and ligands (*26–34*). The seminal paper by Dror et al (*31*) provided insights into the activation mechanism of the *β*_2_AR by monitoring how a crystallized active state conformation relaxed to an inactive state upon removal of the intracellular binding partner, a G protein mimicking nanobody, using MD simulations. Although key conformational changes involved in the transition from active to inactive conformations were identified in these simulations, this approach has inherent limitations. Indeed, understanding the roles of different microswitches in activation and the strength of the coupling between these requires mapping of the relevant free energy landscapes, which are still too costly to calculate using brute-force MD simulations. Enhanced sampling methods, on the other hand, provide a means to explore the conformational landscapes of proteins to a relatively low computational cost (*35*) and have been exploited to study the activation mechanism of GPCRs(*26* –*28, 30, 32* –*34*).

In this work, we aimed to identify the most probable path describing the transition between inactive and active states of *β*_2_AR and to characterize the full conformational ensemble along this pathway at an atomistic level of resolution. A new version of the string method with swarms of trajectories was first developed for this purpose. The free energy landscapes associated with *β*_2_AR activation revealed that whereas agonist binding is only loosely connected to outward movement of TM6, the coupling between microswitch pairs in the transmembrane region ranges from weak to strong. We observed sodium interacting with several binding sites in a state-dependent manner that was consistent with experimental data. Our approach also allowed us to investigate the influence of Asp79^2.50^ protonation and sodium ions on the energy landscape. A protonated (neutral) Asp79^2.50^ favored active-like conformations and led to a distinct conformational state of the NPxxY motif, which closely resemble agonist-bound and active structures of other class A GPCRs. Finally, we demonstrated that the collective variables (CVs) used in this study discriminate active from inactive class A GPCR structures in general. By using the activation path of *β*_2_AR as input to simulations of other receptors, our approach can therefore likely be transferred to study activation of other class A GPCR at a modest computational cost.

## Methods

All swarm of trajectories simulations were initiated with the coordinates from the crystal structure of the active *β*_2_AR in complex with the agonist BI-167107 (from here on referred to as the *agonist*) and an intracellular nanobody (PDB ID 3P0G) (the first two simulation systems in Table S1). The Asn187Glu mutation in 3P0G was reverted and Glu122^3.41^ was protonated due to its localization in a hydrophobic pocket, as has been common practice in other simulations of this receptor (*36*). Residues His172^4.64^ as well as His178^4.70^ were both protonated at the epsilon position. The systems were parametrized using the CHARMM36m force field (*37*) and the TIP3P water model (*38*). The protein was inserted in a POPC (*39*) bilayer and solvated in explicit solvent. Na^+^ and Cl^−^ ions were added at 0.15M concentration. System preparation was performed using CHARMM-GUI (*40*). MD simulations were run with GROMACS 2016.5 (*41*) patched with plumed 2.4.1 (*42*) under a 1 atm pressure and at 310.15 K temperature.

To identify CVs, we performed clustering (*43*) on the frames from an unrestrained MD simulation trajectory of *β*_2_AR (condition A in Dror et al.) (Fig. S1). The CVs were selected by training a multilayer perceptron classifier (*44*) using as input all the inter-residue distances and as output the cluster ID, followed by using Deep Taylor decomposition (*45*) to find key distances that could discriminate between clusters.

The endpoints of the main strings describing the transition between inactive and active states (subscript _*t*_) were fixed to the output coordinates of equilibrated structures of each state (PDB IDs 2RH1 (*5*) and 3P0G (*6*), respectively). The initial path for simulations Holo_*t*_ was guessed using data from Dror et al.: a rough estimate of the free energy landscape was calculated from the probability density landscape estimated using the Scikit-learn (*44*) kernel density estimator with automatic bandwidth detection (Fig. S2). Two additional short strings were set up to increase sampling in the active (subscript _*a*_) and inactive (sub-script _*i*_) regions. The average, partially converged path between iterations 20-30 from Holo_*t*_ was used as input path for simulations Apo_*t*_ and Holo_*Ash*79,*t*_. All active (Apo_*a*_, Holo_*a*_ and Holo_*Ash*79,*a*_) and inactive (Apo_*i*_, Holo_*i*_ and Holo_*Ash*79,*i*_) substrings were initiated as straight paths between the endpoints. Even though the system was initiated from the nanobody bound structure 3P0G, the active state substring sampled conformations closer to the G protein bound structure 3SN6 (Fig. S3) (*7*). The swarms of trajectories simulations with optimizations (Supplementary Methods, Fig. S4 and S5) were run for 300 iterations, at which point the strings had not changed on average for many iterations (Fig. S6 and S7) and posterior distribution of free energy profiles given the data was narrow (Fig. S8).

By discretizing the system into microstates, or bins, it is possible to use the short trajectories from the swarms to create a transition matrix and derive the free energy distribution of the system (*46*) along a given variable (Fig. S9). In practice, the transition probabilities *T*_*ij*_ of the transition matrix *T* can be estimated from the normalized number of transitions, *N*_*ij*_, from bin *i* to bin *j*: *T*_*ij*_ = *N*_*ij*_*/* ∑_*k*_ *N*_*ik*_. The transition matrix of a physical system at equilibrium is constrained by detailed balance, such that for the stationary probability distribution, *ρ*: *ρ*_*i*_*T*_*ij*_ = *ρ*_*j*_*T*_*ji*_. A Metropolis Markov chain Monte Carlo (MCMC) method was used to sample over the posterior distribution of transition matrices, given the unregularized elements of *T*_*ij*_ (*47*), and thereby obtain a distribution of free energy profiles for *ρ* (Fig. S8, S10 and S11). All code to run the simulations and reproduce the results in this paper is available for download (*48*).

## Results

### Optimizing the enhanced sampling protocol for increased efficiency

We aimed to compute the most probable transition pathway linking the inactive and active states of *β*_2_AR and the relative free energy of the states lining this pathway. For this purpose we chose the string of swarms method (*49*). In this framework, the minimum free energy path in an N-dimensional collective variable (CV) space is estimated iteratively from the drift along the local gradient of the free energy landscape (Fig. S5). From each point on the string, a number of short trajectories are launched (a swarm), which enables us to calculate the drift. The string is then updated considering this drift and reparametrized to ensure full sampling of the configurational space along the pathway. Convergence is reached when the string diffuses around an equilibrium position. The method allows to sample a high-dimensional space at a relatively inexpensive computational cost since it only samples along the one-dimensional path of interest.

We characterized the pathway linking equilibrated conformations originating from active and inactive structures of the the *β*_2_AR (PDB IDs 3P0G and 2RH1, respectively), adding two short strings spanning the active and in the inactive regions to increase sampling of the end state environments (see Methods and Supplementary Methods). First, we characterized a CV set that embeds receptor activation by automatically inferring the inter-residue distances that are related to activation. Based on one of Dror et al.’s simulation trajectories of spontaneous deactivation of the *β*_2_AR (*31*), we identified metastable states by clustering simulation configurations, followed by classification of these by training a fully connected neural network to identify states (*50*). The most important input features for classification were identified via deep Taylor decomposition (*45*) and taken as CVs (Fig. 2 and S1). The set of CVs we inferred was a network of interatomic distances between all seven TM helices (Fig. 2). Encouragingly, this five-dimensional CV set captured the main degrees of freedom implicated in *β*_2_AR activation, including the large outward movement of TM6 and a smaller inward shift of TM7 at the intracellular face of the receptor.

**Figure 2:**
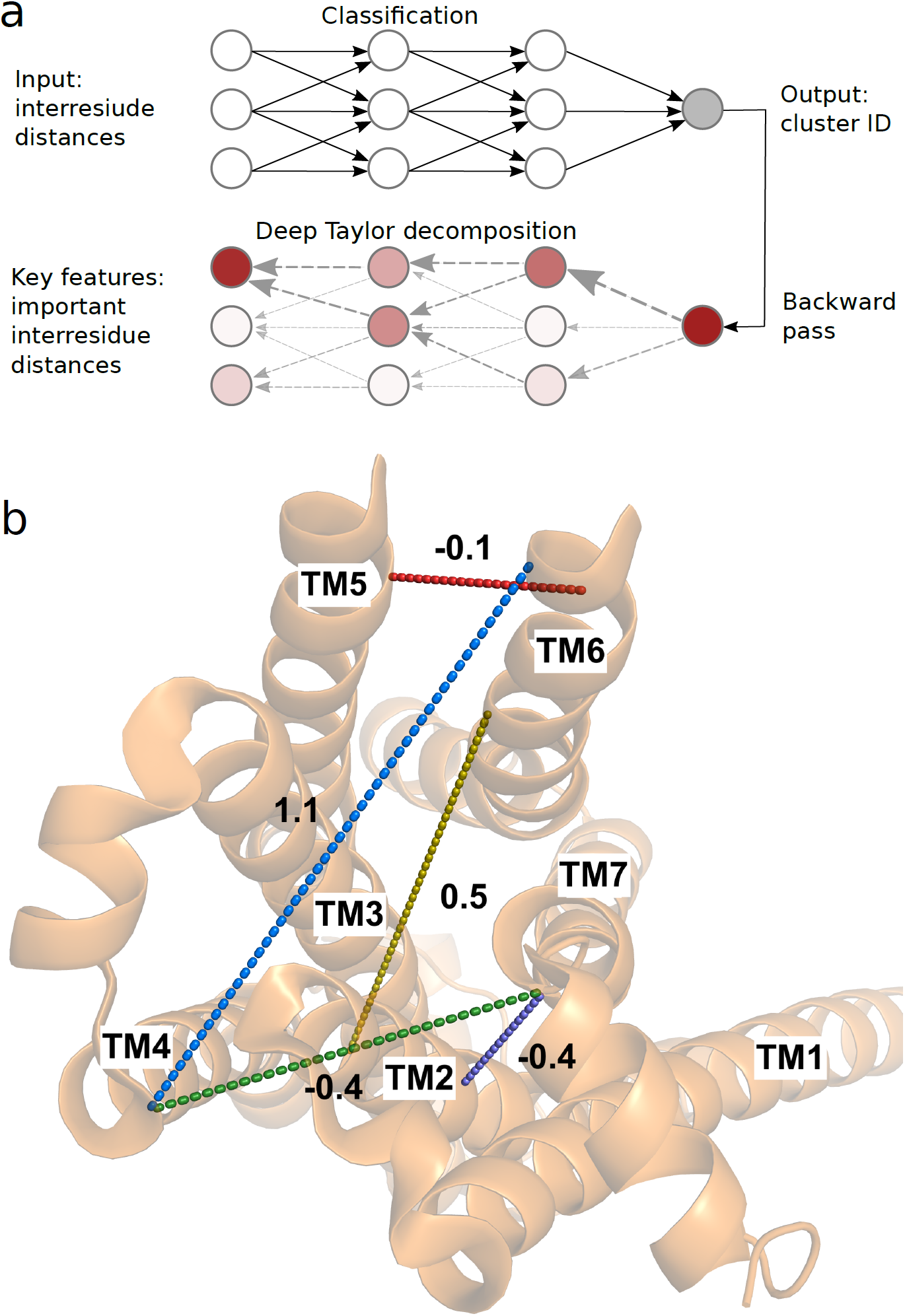
The five-dimensional collective variable (CV) space used to optimize the minimum free energy path was identified in a data-driven manner. (**a**) A fully connected neural network was trained to classify configurations in active, intermediate, and inactive metastable states (clusters). Deep Taylor decomposition was then used to identify the most important input inter-residue distances for the classification decision. The top-ranked distances were used as CVs. (**b**) The five CVs used in this work projected onto an intracellular view of the active crystal structure (PDB ID 3P0G). The CVs corresponding to distances TM2-TM7, TM6-TM4, TM7-TM4, TM3-TM6 and TM6-TM5 defined in Table S2, are shown as purple, blue, green, yellow and red dashed lines, respectively. The change of these distance CVs from the inactive to the active state structures is reported in nanometer (nm).

To speed up convergence of the string optimization, we initiated our string simulation from a rough guess of the minimum free-energy landscape. The latter was obtained by estimating the density of points from Dror et al.’s trajectory in this CV space (Fig. S2). We also introduced algorithmic improvements to the string of swarms method: we adaptively chose the number of trajectories in a swarm, gradually increased the number of points on the string and introduced a reparametrization algorithm that improves performance as well as promotes exchanges of configurations between adjacent string points (Supplementary Methods, Fig. S5, S4). We carried out 300 iterations of string optimization for each system, considering a number of points on the string ranging from 20 to 43 and swarms consisting of 16 to 32 10 ps trajectories. Because we sought to only sample the vicinity of the most probably activation path, derivation of a converged free energy landscape required a mere total of ∼2-3 *μ*s simulation time (Fig. S6, S7, S8, S10 and S11).

### Minimum free energy pathway of *β*_2_AR activation

We derived the most probable transition path (Fig. S6) between active and inactive states of *β*_2_AR in the absence and prescence of an agonist ligand. The swarm trajectories allowed us to compute transitions between discrete states in the vicinity of the most probable transition path (SI methods and Fig. S9) and to derive the associated free energy landscape describing the outward movement of TM6 and inward shift of TM7 upon activation (Fig. 3).

**Figure 3:**
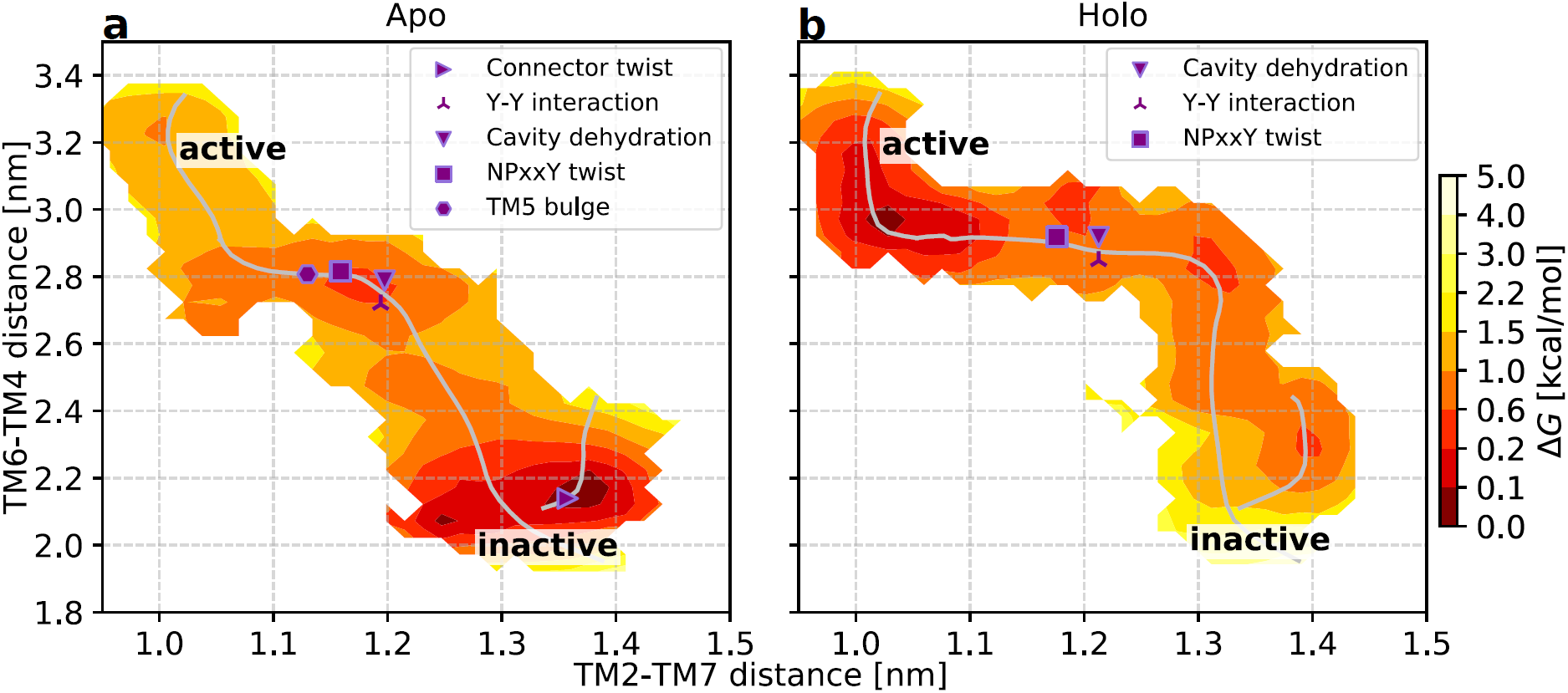
The free energy landscapes are projected onto the first two CVs used to optimize the minimum free energy path. The minimum free energy pathways were optimized in the absence (**a**) and in the presence of the bound agonist (**b**). Characteristic events occurring during activation (see Table S3 for their definition) are located on the string. The active and inactive labels mark the regions close to the active and inactive structures (PDB ids 3P0G(*6*) and 2RH1(*5*), respectively).

For the apo receptor, one distinguishes three minima: one in the active region, one in the inactive one, and an intermediate one between these (Fig. 3a). As anticipated, regions close to the inactive endpoint are stabilized relative to the other two states. Binding of an agonist changes the number of minima to four and shifts the relative stability of states, making regions close to the active conformation of lower free energy (Fig. 3b). A number of characteristic variables (defined in Table S3) were calculated for the last iteration of the swarm of trajectories simulations. By localizing sudden shifts in the values of these parameters, we could pinpoint the location of important events in activation on the string (Fig. 3). In the absence of bound agonist, the connector assumes an active conformation in the early stages of activation. The Asp79^2.50^ cavity then dehydrates, followed by the formation of the Y-Y interaction and NPxxY twist. Finally the TM5 bulge forms before the fully active conformation is reached, confirming an allosteric communication between the orthosteric site and intracellular region.

Binding of the agonist ligand modifies the order in which the helices rearrange, as can be seen when projecting the minimum free energy path along various CVs (Fig. 3 and S6). The most probable activation pathway begins with an outward movement of TM6, followed by the inward shift of TM7. The connector and the bulge in TM5 are locked in their active-like states in the presence of agonist, which enables large outward movements of TM6. Similarly to the apo case, to reach the fully active conformation, the Asp79^2.50^ cavity dehydrates and the Y-Y interaction forms shortly before the NPxxY twists. In the holo simulation, the binding mode of the ligand remained stable along the activation path (Fig. S12a-b; k-l) and it maintained stable interactions with key residues Asp113^3.32^, Ser203^5.43^, Ser207^5.46^ and Asn312^7.38^ (Fig. S12a-j). Occasionally, the closest-heavy atom distance from the ligand to Ser203^5.43^ was almost 4 Å, although these fluctuations were independent of the receptor state (Fig. S12e-f).

### Coupling between orthosteric and G-protein binding sites

As the final configurations of the string are at equilibrium in all dimensions, the trajectories from the last iterations can be used to compute the free energy landscape as a function of any variables (SI methods). This allowed a detailed analysis of how conformational changes induced by agonist binding propagate through the receptor to the G protein binding site. The roles of conserved microswitches were assessed by comparing free energy landscapes in the absence and presence of a bound agonist. We first evaluated how the TM5 bulge, which reflects how the binding site contracts upon activation, influences the intracellular distance between TM6 and TM3 (Fig. 4a,b). In the absence of bound agonist, the receptor accessed both active and inactive conformations of the binding site, with the minimum of the free energy located close to the inactive state distance between TM3 and TM6. The TM3-TM6 distance could be as large as 1.5 nm when the ligand binding site was in the active conformation, an observation compatible with basal activity. Agonist binding led to the stabilization of the TM5 bulge in the orthosteric site (Fig. 4b). Although both inactive and active conformations remained accessible in the presence of the ligand, the minimum of the TM3-TM6 distance was shifted towards a more active-like state. A remarkable long-range allosteric coupling (>2 nm) between the orthosteric and G protein binding sites was hence captured by our simulations. The 0.5 nm outward movement of TM6 at the minimum of the landscape (Fig. 4a,b) is smaller than that observed in active crystal structures, in agreement with experiments demonstrating that the fully active conformation can only be stabilized in the presence of an intracellular partner (*23, 51*).

**Figure 4:**
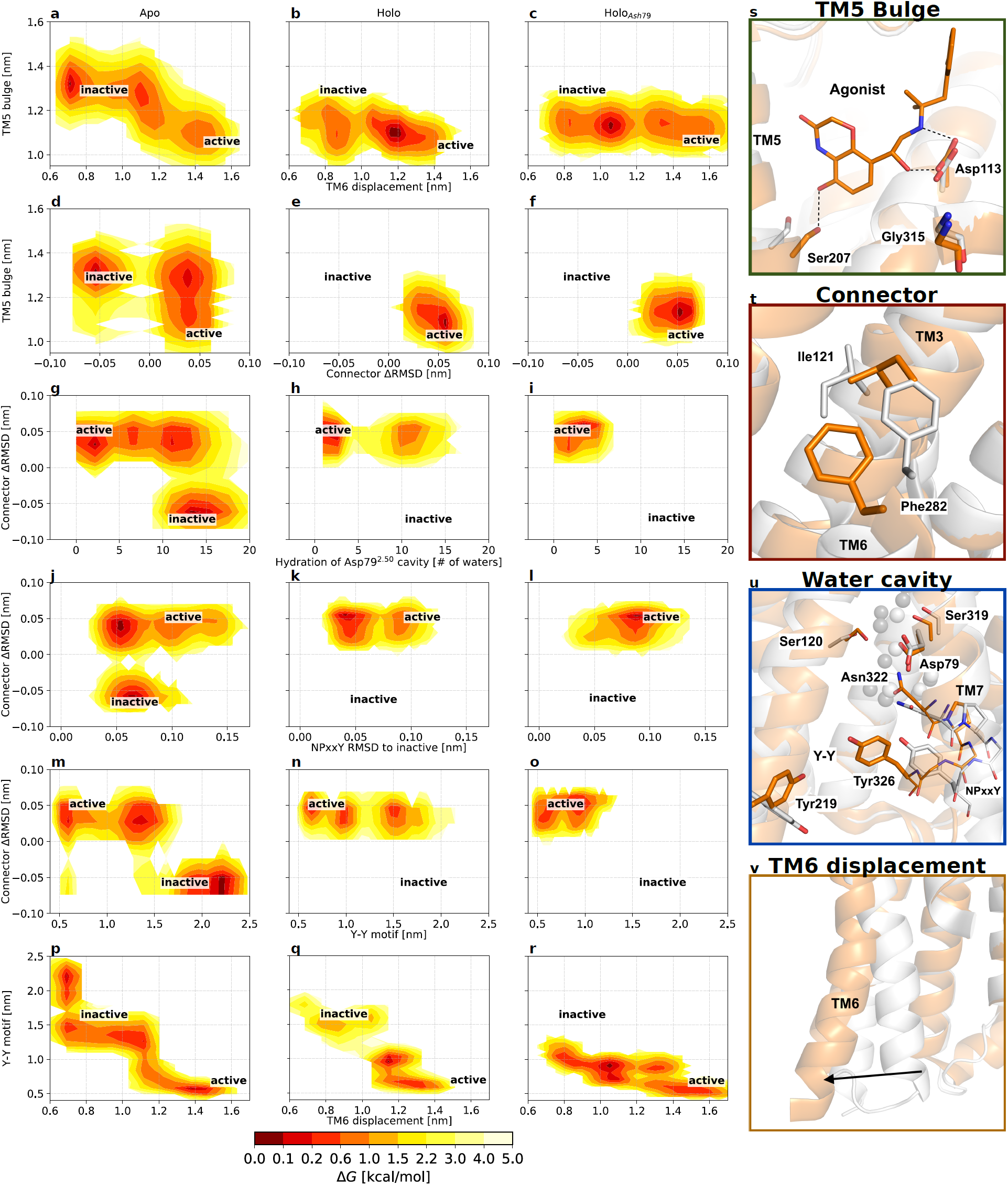
Free energy landscapes projected along variables of interest highlight changes in the pairwise coupling of microswitches following binding of an agonist ligand (middle column) and protonation of conserved residue Asp79^2.50^ (right column). The free energy landscapes are projected along (**a-c**) The TM5 bulge (distance between Ser207^5.46^ and Gly315^7.41^, representing the ligand binding site contraction) and the distance between Leu272^6.34^ and Arg131^3.50^, representing the outward movement of TM6; (**d-f**) The TM5 bulge (distance between Ser207^5.46^ and Gly315^7.41^) and difference between the RMSD of Ile121^3.40^ and Phe282^6.44^ heavy atoms to the active and inactive crystal structures (*5, 6*), representing the connector region ΔRMSD; (**g-i**) the connector region ΔRMSD and the number of water molecules within 0.8 nm from Asp79^2.50^, representing the hydration of the Asp79^2.50^ cavity; (**j-l**) the connector region ΔRMSD and the RMSD of the NPxxY motif relative to the inactive structure 2RH1; (**m-o**) the connector region ΔRMSD and the distance between the two tyrosines implicated in the Y-Y interaction (Y-Y motif). (**p-r**) the Y-Y motif distance and the displacement of TM6. Active and inactive state regions are labeled for each variable pair. Regions of low free energy are shown in red and of high free energy in light yellow. Free energies are reported in kcal/mol. See Table S2 for microswitch definitions. (**s-v**) Vignettes showing the conformation of the different microswitches in the active and inactive structures 3P0G (orange) and 2RH1 (white): (s) the TM5 bulge shifting Ser207^5.46^ inward, (**t**) the connector region containing Ile121^3.40^ and Phe282^6.44^, (**u**) the Asp79^2.50^ cavity, NPxxY motif, and the two tyrosines Tyr219^5.58^ -Tyr326^7.53^ of the Y-Y motif, and (**v**) the outward movement of TM6 upon activation.

### Propagation of activation through microswitches

Structural changes in the orthosteric site of the *β*_2_AR (Fig. 4s) have been proposed to propagate towards the intracellular part via a “connector” centered around Ile121^3.40^ and Phe282^6.44^ (*6*) (Fig. 4t). Whereas we found the TM5 bulge and outward movement of TM6 to be loosely coupled, the free energy landscapes demonstrated that changes in the orthosteric site has a direct influence on the connector region (Fig. 4d,e). In the absence of agonist, both inactive and active conformations of the connector were populated and the connector region could assume an active-like state even if the TM5 bulge was inactive (Fig. 4d). In contrast, agonist binding resulted in a single free energy minimum where both the TM5 bulge and the connector were constrained to their active conformations (Fig. 4e).

From the connector region, activation is propagated via several conserved motifs (Fig. 4g,h,j,k,m,n and u) (*3*). To investigate the communication between microswitches in the core of the TM region, we analyzed if the connector was coupled to solvation of the cavity surrounding Asp79^2.50^, to the conformation of the NPxxY motif, and to the conformation of the Y-Y motif. In the apo state, an inactive connector region was tightly coupled to a hydrated cavity with ∼ 10−17 waters (Fig. 4g) and an inactive NPxxY motif (Fig. 4j). This result is consistent with a high-resolution crystal structure of an inactive *β*_1_ adrenergic receptor (PDB ID 4BVN), in which an ordered solvent network in this region was identified(*17*). In contrast, the active conformation of the connector was compatible with both the fully hydrated cavity and a desolvated state with only four to five water molecules as well as an inactive and active NPxxY motif. Interestingly, the connector configuration was more tightly coupled to the Y-Y motif. An inactive connector was coupled to a broken Y-Y interaction whereas it was in intermediate state or fully formed with an active connector (Fig. 4m).

The free energy landscapes suggested that binding of agonist reduce the number of inactive-like states of the Asp79^2.50^ cavity, NPxxY, Y-Y motifs (Fig. 4h,k,n) as well as the orientation of the Met82^2.53^ (Fig. S11), a residue located one helix turn above Asp79^2.50^ that has been the focus of NMR studies(*52*). For the Asp79^2.50^ cavity and NPxxY motif, both active and inactive conformations were accessible in the presence of agonist, but the active conformations were more favored energetically (Fig. S13). The presence of agonist resulted in an energy landscape with one active-like and two intermediate distances of the Y-Y motif. The two latter minima were shifted by approximately 0.1 nm towards the active conformation compared to the apo state (Fig. 4n).

The final combination of microswitches connected the Y-Y motif to the motion of TM6 (Fig. 4p,q). In the apo state, there was a relatively tight coupling between the Y-Y interaction and the TM3-TM6 distance, with several metastable states lining the minimum free energy pathway between inactive and active conformations (Fig. 4p). Agonist binding did not alter the coupling between these two microswitches, but generally tilted the free energy landscape towards more active states along both of these dimensions (Fig. 4q).

Access to the free energy landscapes in absence and presence of ligand thus enables us to characterize the involvement of each microswitch in the transmission of allosteric coupling between the orthosteric ligand and the intracellular partner binding sites.

### State-dependent sodium binding

Several structures of class A GPCRs solved in the inactive state have revealed a sodium ion bound to the conserved Asp79^2.50^ (Fig. 5a) (*18*). Sodium binding to this residue has also been studied for several class A GPCRs with simulations (*53–55*). To investigate potential interactions between Asp79^2.50^ and sodium ions, which were initially randomly added at physiological concentration to the simulation system, we calculated the free energy landscape as a function of TM6 displacement and the distance between Asp79^2.50^ and the closest sodium ion (Fig. 5b, c and Fig. S11j-l). In the apo form, five meta-stable states were identified (Fig. 5b). In the active-like conformation of TM6, the closest sodium interacted with a specific site in the second extracellular loop. Notably, sodium ions have been confirmed to bind in this pocket in crystal structures of adrenergic receptors (Fig. 5a) (*11*). In an intermediate conformation of TM6, the closest sodium was either bound to the second extracellular loop or descended into the binding site and formed a salt bridge to Asp113^3.32^. Finally, in the completely inactive state of TM6, the closest sodium ion either remained bound to Asp113^3.32^ or was located in the Asp79^2.50^ cavity. Sodium was hence only present in the Asp79^2.50^ cavity when TM6 had completely relaxed to an inactive conformation and even small increases of the TM3-TM6 distance were incompatible with ion binding to this site. In the agonist-bound receptor with Asp113^3.32^ ionized, sodium remained strongly bound to the second extracellular loop irrespective of the TM3-TM6 distance (Fig. 5c). This difference is likely due to that access to the binding site via the extracellular side being blocked by the bound ligand (*19, 53, 55*), but could also be a result of conformational changes induced by the agonist. Spontaneous Na^+^ binding to appropriate protein regions further confirms the relevance of the conformational sampling enabled by our computational protocol.

**Figure 5:**
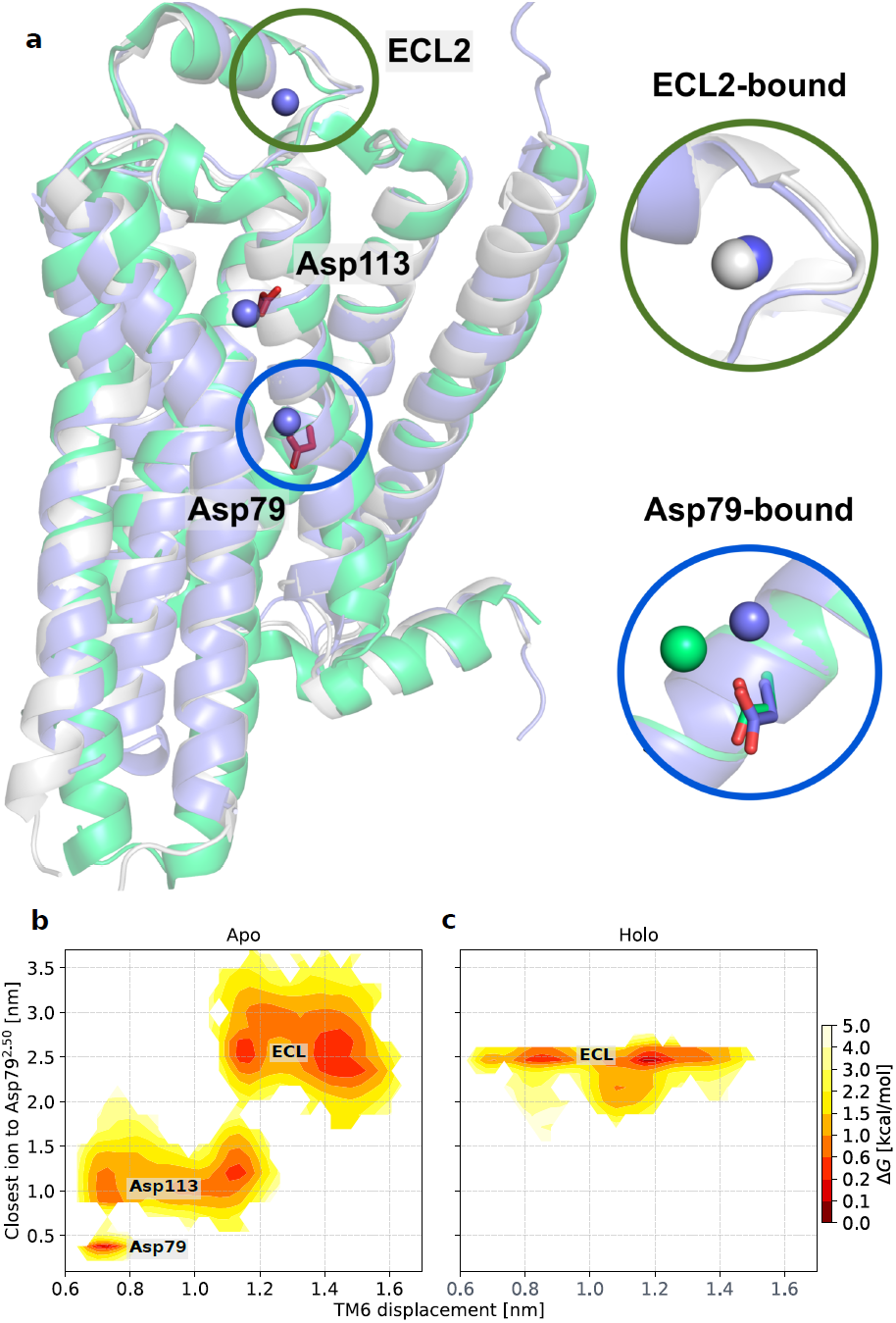
Sodium ions bind to three sites in a state-dependent manner. (**a**) The inactive *β*_1_AR (PDB code: 4BVN (*17*)), *β*_2_AR(PDB code: 4LDE (*11*)) structures and representative inactive state simulation snapshot are shown as green, white and blue ribbons, respectively. Representative positions of sodium ions in the MD simulations are shown as blue spheres. In the insets, a comparison to the positions of sodium ions found in the crystal structures of active *β*_2_AR (white sphere) and inactive *β*_1_AR (white spheres) are shown. (**b, c**) Free energy landscapes along the TM6 displacement and the closest distance between Asp79^2.50^ and a sodium ion for the apo and holo simulations.

### Impact of Asp79^2.50^ protonation

Protonation of two ionizable residues, Asp79^2.50^ and Asp130^3.49^, has been proposed to be involved in GPCR activation (*20, 31, 56*). In particular, MD simulations have indicated that Asp79^2.50^, the most conserved residue among class A GPCRs, has a pK_*a*_ value close to physiological pH and that the ionization state of this residue changes upon activation (*19, 20*). As a previous simulation study of the *β*_2_AR showed that Asp79^2.50^ (but not Asp130^3.49^) may alter the activation pathway(*31*), we repeated the calculations of the minimum free energy pathway of activation with this residue in its protonated (neutral) form (Fig. S7).

The free energy landscapes describing changes in the orthosteric site were similar to those obtained in simulations with Asp79^2.50^ ionized (Fig. 4c,f). There was a weak coupling between the TM5 bulge and the intracellular region, with two major energy wells describing the conformation of TM6. Compared to the agonist-bound receptor with Asp79^2.50^ ionized, the minima were shifted further towards active-like conformations for the protonated state (Fig. 4c). The TM5 bulge remained strongly coupled to conformational changes observed in the connector region irrespective of the ionization state of Asp79^2.50^ (Fig. 4e,f). The largest effects of Asp79^2.50^ protonation were observed for the hydrated cavity surrounding this residue (Fig. 4h,i), NPxxY (Fig. 4k,l), and Y-Y motifs (Fig. 4n,o): whereas the free energy landscapes showed that both active- and inactive-like conformations of these switches were populated in simulations with ionized Asp79^2.50^, the protonated state only resulted in energy wells close to the active conformation (Fig. 4i,l,o). It was also evident that TM6 was stabilized in more active-like conformations by the protonated Asp79^2.50^ (Fig. 4r). Interestingly, intermediate states in which the Y-Y motif, as well as the NPxxY, adopted an active-like conformation and a small TM6 displacement were also populated (Fig. 4r and S13). Such an intermediate state has been observed, albeit rarely, in simulations considering a protonated Asp79^2.50^ (*31*).

Comparison of representative structures from the simulations of ionized and protonated Asp79^2.50^ in active-like states revealed that two distinct conformations of the NPxxY motif were obtained (Fig. 6). The simulations carried out with ionized Asp79^2.50^ favored structures that were more similar to the crystal structure of the active *β*_2_AR. An alternative conformation of the active NPxxY motif appeared for the protonated Asp79^2.50^, which was not favored energetically in the simulations of the ionized form (Fig. S10k,l and S13) Although this conformation of the NPxxY motif did not match any *β*_2_AR crystal structure, it was strikingly similar to conformations observed in crystal structures of other class A GPCRs in either agonist-bound (serotonin 5-HT_2*B*_ and A_2*A*_ adenosine receptors) or active (angiotensin II type 1) conformations (Fig. 6) (*57, 58*). Our protocol thus allowed us to sample metastable states not yet resolved by experimental structure determination that may play a role in activation.

**Figure 6:**
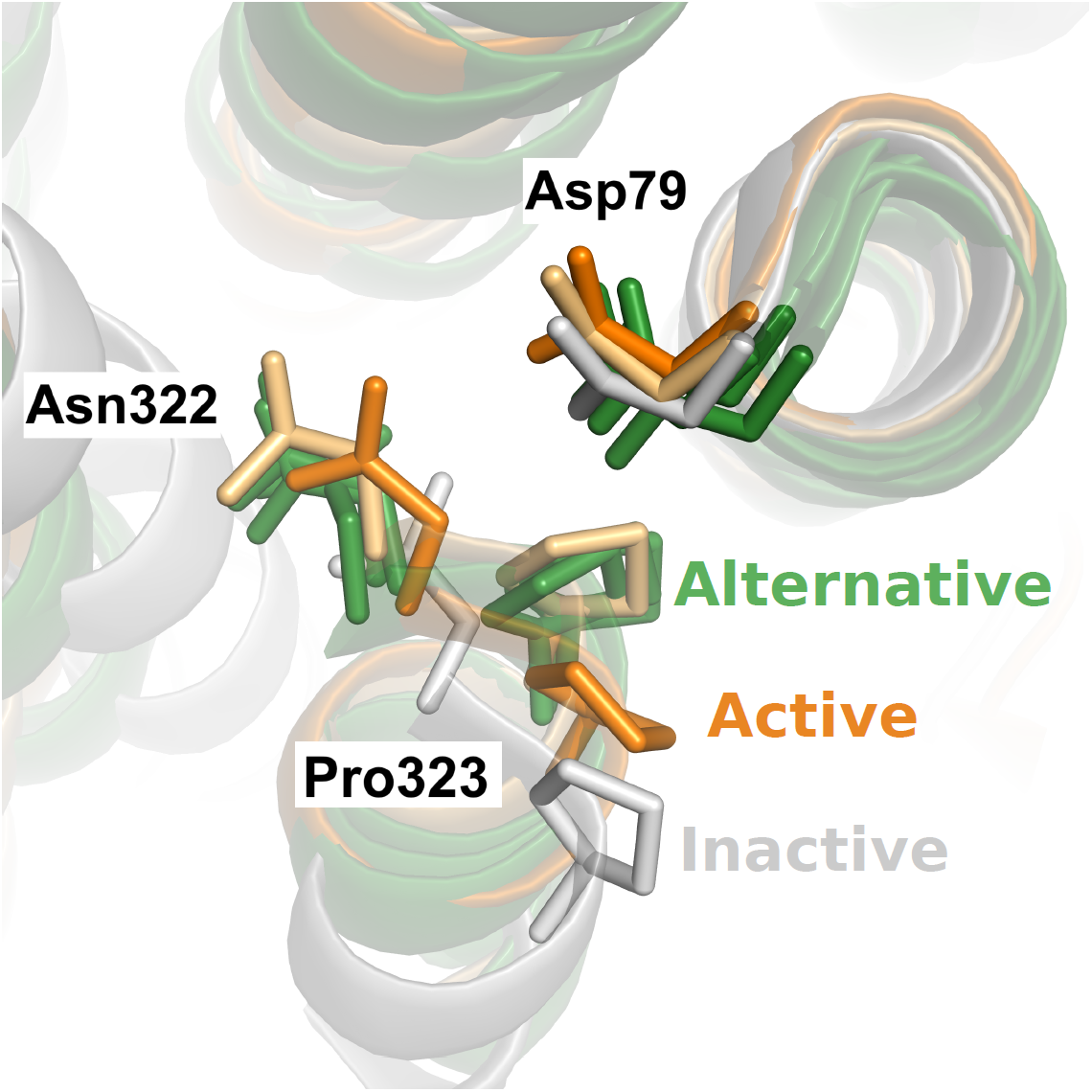
An alternative conformation of the NPxxY was identified along the most probable pathway calculated with a protonated Asp79^2.50^. Representative simulation snapshots of active-like states of the *β*_2_AR with Asp79^2.50^ ionized and protonated are shown in orange and light-orange, respectively. Representative simulation snapshots of the inactive-like state of the *β*_2_AR is shown in white. The structural comparison highlights the resemblance between an alternative conformation of the NPxxY motif (light orange) that is favored by Asp79^2.50^ protonation in the simulations and observed in three crystal structures of other class A GPCRs. Crystal structures of the other GPCRs are shown in green (PDB IDs: 6DRX (*57*), Agonist-bound 5-HT_2*B*_ serotonin receptor; 3QAK (*59*), Agonist-bound A_2*A*_ adenosine receptor; and 6DO1 (*60*), Angiotensin II type 1 receptor in an active conformation).

### Identified CVs capture activation of many class A GPCRs

Efficient characterization of free energy landscapes with the string method relies on selection of appropriate CVs, which is a non-trivial task. Here, CVs were derived from a conventional MD trajectory of *β*_2_AR deactivation in a data-driven fashion. Considering that the conformational changes involved in class A GPCR activation are largely conserved (*3*), we explored the possibility of transferring the CVs to the conformational sampling of other GPCRs. We mapped the CVs identified for *β*_2_AR to ten other class A GPCRs with active and inactive structures available (Fig. 7). Strikingly, the active and inactive structures clearly separate in two distinct clusters. Together with the fact that we could capture relevant receptor conformations that were not observed in the MD trajectory used to identify CVs (Fig. 6), this indicates that our approach could potentially be used to characterize the activation of many class A GPCRs.

**Figure 7:**
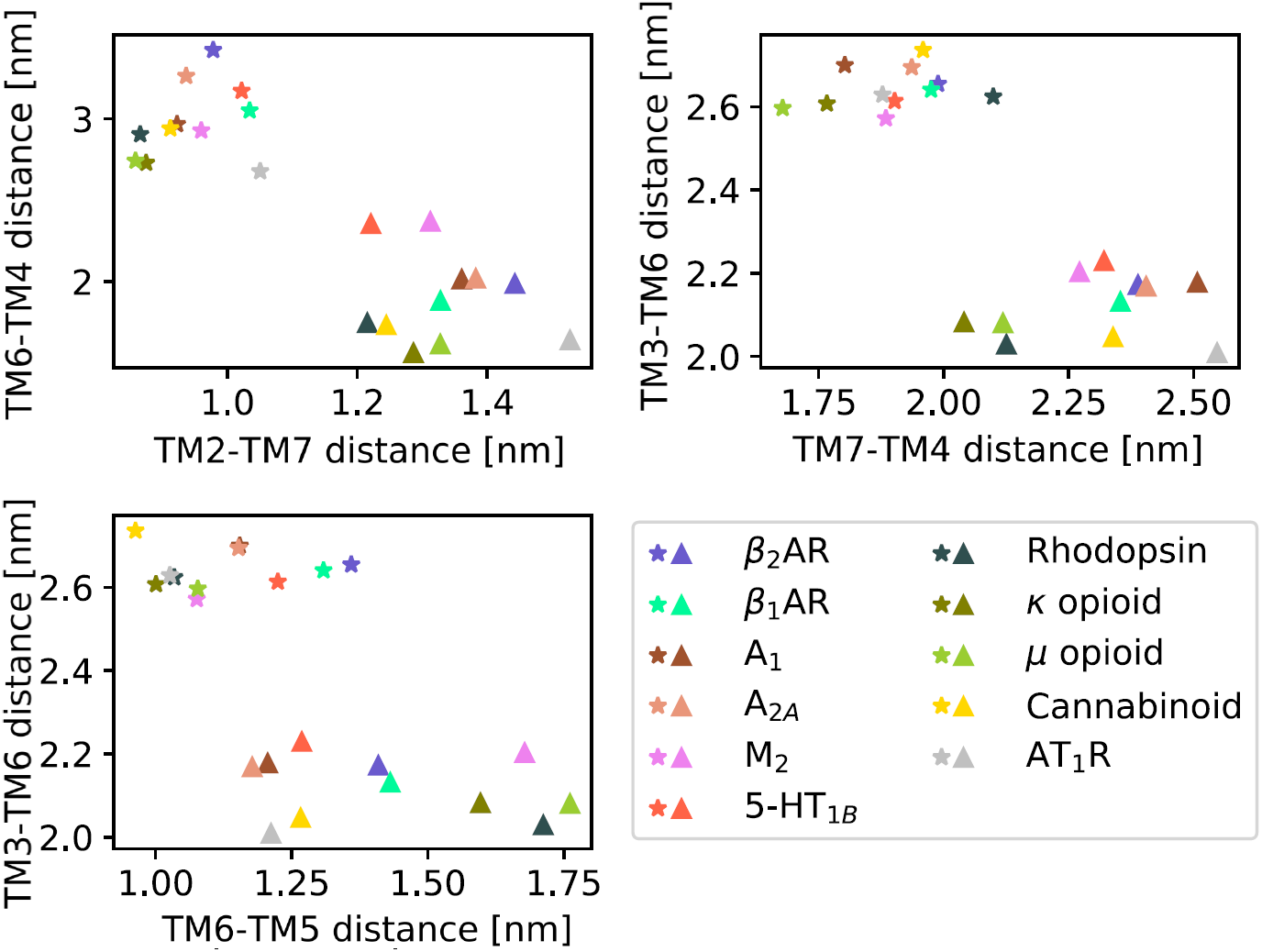
Active and inactive class A GPCR structures cluster into two distinct groups in the 5-dimensional CV space used in this work. *β*_2_AR and ten active (stars) and inactive (triangles) structures of other class A GPCRs are projected onto the five original CVs. Details of the mapping can be found in Table S4. Their clustering into two groups highlights that the activation mechanism of all class A GPCRs can likely be described by the CVs identified in this work.

## Discussion

Experimental structures of the *β*_2_AR in inactive and active conformations provide a basis for molecular understanding of GPCR signaling. However, it has become increasingly clear that the static structures do not capture all functionally relevant states involved in activation of these molecular machines (*26, 27, 32, 33*). In a pioneering study, Dror et al. gained insights into the activation pathway of the *β*_2_AR from long-timescale MD simulations (*31*). Despite this computational tour-de-force enabled by the development of hardware specialized for MD, these simulations did not allow to quantify the accessibility of different conformational states. The approach proposed in this work builds on the data generated by Dror et al. and on methodologies suggested in previous molecular simulation studies (*27, 29, 30, 33*) to further allow to assess the impact of agonist binding on microswitches involved in activation.

Our work recapitulates a number of key findings of previous enhanced sampling (*27, 29, 30, 32, 33*) and long-timescale scale simulations of GPCRs (*31*). In agreement with previous work, our simulations revealed a complex conformational landscape of the *β*_2_AR with only weak coupling between the othosteric site and G protein binding site. Through the analysis of the free energy landscapes we could quantify the coupling between spatially connected microswitches in the transmembrane region, which ranged from strong to relatively weak and was influenced by protonation of Asp79^2.50^. Our study also further illustrates that the energy landscapes depend on the variables chosen to project the conformational states. This is not an artifact of the protocol but rather an inherent limitation of dimensionality reduction. In particular, it is clear that considering only three states along the activation path (an active, intermediate and inactive state) does not allow to capture the complexity of the conformational changes induced by ligand binding (*26, 27, 32, 33*). The present work, through the resolution of the complete conformational ensemble lining the most probable activation pathway, can aid in determining the placement of spectroscopic probes to monitor the distribution of specific microswitches under different conditions.

Our work shows agreement with spectroscopy studies although a quantitative comparison remains challenging: whereas the *β*_2_AR crystallized in complex with agonists in the absence of intracellular partner (e.g. G protein or G protein-mimicking nanobody) are similar to identical to those determined with antagonists (*36*), NMR spectroscopy experiments have demonstrated that agonist binding stabilizes other conformations in certain parts of the TM region, e.g. close to Met215^5.54^ and Met82^2.53^ (*52*). For example, the region surrounding Met82^2.53^, located one helical turn above Asp79^2.50^, was shown to adopt three conformations in the prescence of a neutral antagonist (with similar signaling properties as the apo receptor). A bound agonist, on the other hand, restricted this region to a single active-like state (*51, 52*). Similar to these experiments, our free energy landscapes demonstrate that several conformations of the connector, Asp79^2.50^ cavity and the orientation of Met82^2.53^ are available in the apo condition (Fig. S11d-f and S10g-i,m-o). Especially, Fig. S11d-f shows three conformational Met82^2.53^ states of comparable free energy for the apo receptor, whereas the landscape is significantly shifted to favour a single active state conformation for the agonist bound receptor. Protonation of Asp79^2.50^ further increases this shift. In the presence of agonist, the connector is locked in a single state and a desolvated state of the Asp79^2.50^ cavity is stabilized, which creates a more active-like receptor conformation in the vicinity of Met82^2.53^. In agreement with the NMR data, we also find that the agonist cannot stabilize the fully active conformation of the receptor and that TM6 accesses several intermediate conformations that are distinct from those observed in crystal structures.

The rapidly increasing number of crystal structures provides insights into how agonist binding to GPCRs, despite recognizing disparate molecules, can result in very similar conformational changes. The bulge of TM5, which we found to be strongly coupled to the connector region, is is also present in active-state structures of other GPCRs (*61, 62*). Whereas this conformational change is the result of a direct interaction with TM5 in the case of the *β*_2_AR, agonist interactions with other helices appear to indirectly influence the same region in the case of the *μ* opioid and A_2*A*_ adenosine receptor (*61, 62*). In other cases, the connector region was found to have a similar arrangement in the active and inactive state (e.g. the M_2_ muscarinic receptor) (*63*). In such cases, receptor activation may be controlled by direct modulation of the Asp79^2.50^ cavity, e.g. the via the conserved Trp286^6.48^. Several recent experimental studies have demonstrated that Asp79^2.50^ and residues forming the hydrated TM cavity play an important role in signaling and can even steer activation via G protein-dependent and G protein-independent pathways (*25, 64, 65*). One mechanism by which Asp79^2.50^ could control the receptor conformation is via its protonation state. Agonist binding destabilizes the water network in the solvated TM cavity, which may lead to a larger population of protonated Asp79^2.50^ and disrupt binding of sodium to this pocket (*19, 20*). In turn, the protonated Asp79^2.50^ stabilizes a structure of the NPxxY that has been observed for other class A GPCRs crystallized in active and active-like states, suggesting that this alternative conformation of TM7 may be relevant for function. NMR experiments have shown that different ligands stabilize different active states(*66*). In particular, agonists that preferentially signal via arrestin mainly affect the conformation of TM7, whereas a neutral agonist stabilizes both the G protein as well as the arrestin signaling state. While two distinct active-state conformations of the NPxxY motif were identified in our simulations for a neutral agonist, a ligand with different biased signalling properties may preferentially stabilize one of these states.

Interestingly, the ionized and protonated forms of Asp79^2.50^ also stabilize different TM6 conformations, which could change the intracellular interface that interacts with G proteins and arrestins(*9*). These results, combined with the fact Asp79^2.50^ is the most conserved residue in the class A receptor family, support that this region is a central hub for controlling class A receptor activation.

With the protocol developed herein, we sampled enough transitions along the activation path to obtain free energy profiles of GPCR activation by accumulating a few microseconds of total simulation time. Compared to regular MD simulations, the optimised string of swarms method can thus provide reliable energetic insights using 1-2 orders magnitude less simulation time than plain MD simulations (*31, 32*), and comparable to other enhanced sampling methods such as GaMD (*26*). From a practical point of view, the short trajectories in the swarms of trajectories method are easy to run in parallel with minimal communication overhead even in a heterogeneous computational environment. An important consideration that has guided our choice of enhanced sampling methodology is that the method has the advantage to function well in high dimensional space, i.e. with many CVs. This is because we only optimize the one-dimensional string, instead of opting to sample the entire landscape spanned by the CVs. This means we can utilize a high-dimensional CV space, thus alleviating the need to reduce the dimensionality of the conformational landscape to 1 or 2 dimensions, as is done in most CV-based methods such as umbrella sampling or metadynamics (*67*).

On the other hand, a well-known limitation of the string with swarms method is that it only guarantees to converge to the most probable path closest to the initial guess of the path, and not necessarily the globally most probable path. The naive assumption of a straight initial path is not guaranteed to converge to the latter. Here we have proposed to alleviate this shortcoming by exploiting previous knowledge of the activation pathway and deriving an initial guess of the pathway likely to be close to the globally most probable path. An initial pathway can also be transferred from a similar system (as revealed in Fig. 7) or inferred from available experimental data. If multiple pathways are nevertheless expected, the protocol presented herein provides the tools necessary to compare them: the swarms from separate string simulations can be included in the same transition matrix and be used to compute a single free energy landscape. It is also worth noting that the relative free energies of the endpoint conformations (evaluated by integration over the endpoints basins) do not depend on the transition pathway and should be estimated correctly. The protocol is also applicable to complex transitions involving many intermediate states: in such case, one may launch multiple strings to explore different parts of the activation path, and let every substring converge separately, eventually combining the transitions derived from them to yield a single free energy landscape. Finally, we note that the string of swarms method can also be used as a complementary method to instantiate Markov State Models (MSM) simulations (*68*).

Despite the major progress in structural biology for GPCRs, many aspects of receptor function remain misunderstood. Insights from atomistic MD simulations will continue to be valuable tools for interpreting experimental data. We expect our methodology to allow further insights into how binding of different ligands influences the conformational landscape, potentially making it possible to design ligands with biased signaling properties (*25*). The method is equally well suited to study the effect of allosteric modulators and the influence of different lipidic environments (*69*). As the same approach can likely be transferred to other class A GPCRs, future applications will shed light on the common principles of activation as well as on the details that give each receptor a unique signaling profile, paving the way to the design of more effective drugs.

## Abbreviations

MD: Molecular Dynamics
GPCR: G protein-coupled receptor
CV: Collective Variable
*β*_2_AR: *β*_2_ adrenergic receptor

## Acknowledgement

This work was supported by grants from the Gustafsson Foundation and Science for Life Laboratory to JC and LD. The work was also supported by grants from the Swedish Research Council (2017-4676) and the Swedish strategic research program eSSENCE to JC. The simulations were performed on resources provided by the Swedish National Infrastructure for Computing (SNIC) at PDC Centre for High Performance Computing (PDC-HPC). The authors thank D.E. Shaw Research for providing access to their MD trajectories.

## Supporting Information Available

Supplementary Tables S1-S4, Supplementary Methods (System Preparation, MD simulation details, CV selection, Swarms of trajectories simulation setup, Algorithmic improvements to the swarms of trajectories method and Free Energy Computation) and Supplementary Figures S1-S13.

This material is available free of charge via the Internet at http://pubs.acs.org/.

