## Supplementary Information for "Energy landscapes reveal agonist control of GPCR activation via microswitches"

#### Running header

Energy landscapes reveal agonist control of GPCR activation via microswitches

### Supplementary Tables

Table S1: Simulation systems equilibrated to define string endpoints

| Condition | PDB structure | # atoms | Used for string simulation trajectories |
|---|---|---|---|
| apo | 3P0G (active) | 71734 | yes |
| holo | 3P0G (active) | 68316 | yes |
| apo | 2RH1 (inactive) | 54478 | no, only used to get CV coordinates of endpoints |
| holo | 3SN6 (active) | 74452 | no, only used to get CV coordinates of endpoints |

Table S2: Original CVs and other variables used to characterize activation events or to generate free energy plots

| Name | Definition |
|---|---|
| TM2-TM7 distance | Ala76$^{2.47}$-Cys327$^{7.54}$ C$\alpha$ distance (original CV) |
| TM6-TM4 distance | Glu268$^{6.30}$-Lys147$^{4.39}$ C$\alpha$ distance (original CV) |
| TM7-TM4 distance | Cys327$^{7.54}$-Lys147$^{4.39}$ C$\alpha$ distance (original CV) |
| TM3-TM6 distance | Leu115$^{3.34}$-Leu275$^{6.37}$ C$\alpha$ distance (original CV) |
| TM6-TM5 distance | Lys267$^{6.29}$-Phe223$^{5.62}$ C$\alpha$ distance (original CV) |
| TM6 displacement | Leu272$^{6.34}$-Arg131$^{3.50}$ C$\alpha$ distance |
| TM5 bulge | Ser207$^{5.46}$ and Gly315$^{7.41}$ closest heavy atom distance |
| Connector $\Delta$RMSD | Difference between the average Root-mean-square deviation (RMSD) of Ile121$^{3.40}$ and Phe282$^{6.44}$ heavy atoms to the inactive and active structures 2RH1 and 3P0G |
| NPxxY RMSD to inactive | Residue 322 to 327 RMSD to the inactive structure 2RH1 |
| Hydration of Asp79$^{2.50}$ cavity | Number of water molecules within 0.8 nm of Asp79$^{2.50}$. (distance between water molecules' oxygen atom and residue's closest heavy atom were considered) |
| Closest ion to Asp79$^{2.50}$ | Distance to closest sodium from any Asp79$^{2.50}$ heavy atom |
| Y-Y motif | Tyr219$^{5.58}$-Tyr326$^{7.53}$ C$\zeta$ distance |
| \|Asp79-Asn322\| | Asp79$^{2.50}$-Asn322$^{7.49}$ closest heavy atom distance |
| Met82 $\Delta$RMSD | Difference between the RMSD of Met82$^{2.53}$ heavy atoms to the inactive and active structures 2RH1 and 3P0G |
| Trp286 $\Delta$RMSD | Difference between the RMSD of Trp286$^{6.48}$ heavy atoms to the inactive and active structures 2RH1 and 3P0G |

Table S3: Characteristic events for variables defined in Table S2.

| Name | Definition |
|---|---|
| Connector twist | Connector $\Delta$ RMSD $\leq 0$ nm |
| Cavity dehydration | Less than 5 water molecules within 0.6 nm of Asp79$^{2.50}$ |
| Y-Y interaction | Y-Y motif distance $\leq 0.75$ nm |
| NPxxY twist | NPxxY RMSD to inactive $\geq 0.07$ nm |
| TM5 bulge | Orthosteric site TM5 contraction $\geq 1.2$ nm |

Table S4: Structure-based CV mapping between class A GPCRs with an active and an inactive state PDB structure

| Name | active /inactive | $CV_1$ $\|2.47 - 7.54\|$ | $CV_2$ $\|6.30 - 4.39\|$ | $CV_3$ $\|7.54 - 4.39\|$ | $CV_4$ $\|3.34 - 6.37\|$ | $CV_5$ $\|6.29 - 5.62\|$ |
|---|---|---|---|---|---|---|
| $\beta_2$AR | 3SN6 (*1*) /2RH1 (*2*) | A76-C327 | E268-K147 | C327-K147 | L115-L275 | K267-F223 |
| $\beta_1$AR | 6H7N* (*3*) /5A8E (*4*) | A84-C344 | E285-R155 | C344-R155 | L123-L292 | R284-Y231 |
| $A_1$ | 6D9H (*5*) /5UEN (*6*) | A52-A289 | E229-P121 | A289-P121 | I89-L236 | K228-F204 |
| $A_{2A}$ | 5G53 (*7*) /4EIY (*8*) | A49-A289 | E228-G118 | A289-G118 | V86-L235 | K227-F201 |
| $M_2$ | 4MQt* (*9*) /3UON (*10*) | A66-A441 | E382-T137 | A441-T137 | V105-I389 | R381-S210 |
| Rhod-opsin | 6CMO (*11*) /1F88 (*12*) | A80-I307 | E247-E150 | I307-E150 | L119-V254 | A246-V227 |
| $\kappa$ opioid | 6B73* (*13*) /4DJH (*14*) | A102-A331 | L269-P172 | A331-P172 | Y140-V276 | N268-I250 |
| $\mu$ opioid | 5C1M* (*15*) /4DKL (*16*) | A111-A337 | L275-P181 | A337-P181 | Y149-V282 | N274-I256 |
| Canna-binoid | 6N4B (*17*) /5U09 (*18*) | A160-A398 | D338-R230 | A398-R230 | T197-L345 | M337-L298 |
| 5-$HT_{1B}$ | 6G79 (*19*) /5V54 (*20*) | A92-T370 | E309-P163 | T370-P163 | T131-L316 | R308-Y232 |
| $AT_1$R | 6DO1* (*21*) /4YAY (*22*) | A71-G303 | N235-M142 | G303-M142 | F110-I242 | N235-W219 |

Active structures marked with an asterix (*) are bound to a nanobody.

### Supplementary Methods

#### System Preparation

Four systems were equilibrated and used to define the endpoints for the string simulations, see Table S1. The initial coordinates were taken from PDB structures 3P0G (*23*), 2RH1 (*2*) and 3SN6 (*1*). The intracellular binding partners, if any, were removed. Residues Asp130$^{3.49}$ and His172$^{4.64}$ were not protonated, the Asn187Glu mutation in 3P0G was reverted, Glu122$^{3.41}$ was protonated and residues His172$^{4.64}$ as well as His178 were protonated on the epsilon position. The agonist was either bound in the ligand binding site, referred to as the *holo* condition, or absent from the system, referred to as the *apo* condition. For each system, the protein was inserted in a POPC (*24*) bilayer and solvated in explicit solvent. $Na^+$ and $Cl^-$ ions were added to neutralize the systems' total charge. System preparation was performed using CHARMM-GUI (*25*).

#### MD simulation details

The systems were parametrized using the CHARMM36m force field (*26*) and TIP3P water model (*27*). The ligand had its amine group protonated and was parameterized with CHARMM's general force field using CHARMM-GUI's ligand modeler (*28*).

MD simulations were run with GROMACS 2016.5 (*29*) patched with plumed 2.4.1 (*30*) in an NPT ensemble at 310.15 K. The simulation systems were minimized, thermalized and equilibrated under gradually decreasing constraints, using CHARMM-GUI's default configuration. The time step considered was 2 fs and hydrogen bond lengths were constrained with the LINCS algorithm. Thermostatting was performed using Nose-Hoover thermostats with a time constant $\tau = 1$ ps for protein, membrane and electrolyte while barostatting involved semiisotropic Parrinello-Rahman pressure coupling with a time constant $\tau_p = 5$ ps and compressibility of 4.5 $10^{-5}$ $\text{bar}^{-1}$.

Structural visualizations were made in PyMOL (*31*).

#### CV selection

The collective variables (CVs) used to optimize the minimum free energy pathway should capture slow degrees of freedom of the system, separate distinct functional states and characterize the overall process of activation. In contrast to other enhanced sampling methods where the simulation performance is reduced by increasing the number of CVs, the string is one-dimensional and the computational complexity and expected simulation time of the swarms of trajectories method does not increase with the number of CVs. We thus designed a protocol that allows to derive a relatively low dimensional set of CVs but that does not restrict itself to an artificially low number.

The CVs were derived from an unrestrained MD trajectory of holo $\beta_2$AR (Condition A in (*32*)). The CVs were set to be certain interatomic distances using the following protocol: first, to determine the number of conformational states (clusters) present in the simulation in an unbiased way, we clustered the simulation snapshot using spectral clustering on inverse inter-residue distances and computed the SPECTRUS quality score (*33*, *34*) for a number of clusters ranging from 2 to 12 (Fig S1). The highest quality score was obtained for 3 clusters (Fig S1a). By projecting the cluster onto characteristic variables (*1*, *23*) (TM6-TM3 displacement and the NPxxY twist), the clusters were shown to correspond to the active, the inactive as well as the intermediate state (Fig. S1b) previously identified by Dror et al (*32*). Next, we trained a multilayer perceptron classifier (*35*) using as input all the interresidue distances and as output the cluster ID. The neural network used ReLu activation and was trained using a lbfgs solver [1]. The classifier performed well with less than 1% false classifications during 2-fold cross validation. The classifier was then trained with all available data and finally deep Taylor decomposition (*36*) was used to identify interresidue distances of relevance for the classification. The top five interatomic distances were used as CVs and can be seen in Fig. 2.

The CVs were scaled to generate unitless values ranging from 0 to 1. This is important

[1] Unless otherwise specified, the default settings of scikit-learn were used.

when we compute the drift and reparametrize the string, otherwise CVs corresponding to small-scale movements will be suppressed during reparametrization.

Although it is easy to come up with variations to the scheme presented here for CV discovery, the basic idea is to find human-interpretable CVs which well parametrize the transition between states along the activation path.

There is no requirement that the MD trajectories have to come from unrestrained long-timescale simulations, even though that was used in this paper. Data to use for clustering and CV set parametrization may be obtained by running MD simulations by, for example, removing the ligand to reduce the timescales of deactivation from the active states, mutating a residue, adding a biased force to the system, launching simulations from different crystal structures, increasing the temperature to overcome ergodicic barriers or any other methods which approximately samples states along the activation path.

#### Swarms of trajectories simulation setup

In the string with swarms of trajectories method, the most probable transition path is estimated iteratively from the drift along the local gradient of the free energy landscape. From each point on the string, a number of short trajectories are launched (the swarm), which enables to calculate the drift. The string is then updated considering this drift and reparametrized to ensure full sampling of the configurational space (Fig. S4). Convergence is reached when the string diffuses around an equilibrium position.

All swarms of trajectories simulations were instantiated with the coordinates from the 3P0G structure (the first two simulation systems in Table S1). The endpoints of the main strings describing the transition between inactive and active states (subscript $_t$) were fixed to the output coordinates of the equilibrated structures of 2RH1 and 3P0G. Two additional short strings were setup to increase sampling in the active (subscript $_a$) and inactive (subscript $_i$) regions. The endpoints of the active substrings were set to the output coordinates of the equilibrated structures of 3P0G and 3SN6, while the endpoints of the inactive strings were fixed to the average CV coordinates of the second half of two 20 ns unrestrained trajectories which were launched from the output coordinates of the $\text{Holo}_t$ and $\text{Apo}_t$ MD simulations steered towards 2RH1.

The path for simulations $\text{Holo}_t$ was inferred using the same data as was used for clustering. A rough estimate of the free energy landscape was calculated from the probability density landscape estimated using the Scikit-learn (*35*) kernel density estimator (KDE) with automatic bandwidth detection. The initial pathway was then estimated by letting an initial string iteratively follow the gradient of the free energy landscape estimate until convergence to a pathway where the gradient is approximately zero in any direction perpendicular to the string (Fig. S2). The average, partially converged path between iterations 20-30 from $\text{Holo}_t$ was used as input path for simulations $\text{Apo}_t$ and $\text{Holo}_{Ash79,t}$. All active and inactive substrings for Apo and Holo were instantiated as straight paths between the endpoints. Finally the average values of $\text{Holo}_a$ and $\text{Holo}_i$ for iterations 130-230 used as input to $\text{Holo}_{Ash79,a}$ and $\text{Holo}_{Ash79,i}$ respectively.

The number of points on the strings were incremented at fixed intervals (every 8 iterations) to gradually increase the resolution of the activation paths. For the transition strings, we started with 20 points and added additional points until the string consisted of 43 points. For $\text{Holo}_a$, $\text{Apo}_a$, we started with 4 points and additional points were added until the string reached a length of 14 points. The inactive substrings, $\text{Holo}_i$, $\text{Apo}_i$, started with 4 points and reached a final length of 12 points. $\text{Holo}_{Ash79,a}$ and $\text{Holo}_{Ash79,i}$ both started at 6 points and reached a final length of 12 points. The endpoints were fixed for all strings and not updated between iterations.

The strings were initialized by running steered MD along the initial path with a harmonic restraint potential acting on all 5 CVs (*30*) ($\kappa = 10000$ kJ) gradually moving along the points of the string. The simulation frames with CV value closest to the string points were extracted and equilibrated with a 10000 kJ nm$^{-2}$ harmonic force potential acting on the 5 CVs for 4 ns before the first swarm trajectories of 10 ps each were launched. For later iterations, the trajectories were restrained to the points of the string with a the same force potential for 30 ps per iteration.

The swarms were launched in parallel batches of 4 simulations per node using the GROMACS (*29*) '-multi' option starting from the output checkpoint file of the restrained simulation. The number of batches were set adaptively for every point (see details below) to minimize computational cost without compromising the accuracy of the estimated drift by using a new convergence metric (detailed below). After the first 4 batches, the convergence metric was computed and new swarms were launched if necessary. The convergence limit of the drift was initially set to 0.2.

#### Algorithmic improvements to the swarms of trajectories method

##### Adaptive number of swarm trajectories

The most probable transition path is estimated from the drift along the local gradient of the free energy landscape. Increasing the number of trajectories will reduce the standard error in the estimate but also incur an additional computational cost. Thus we suggest an adaptive approach for choosing the number of swarms (Fig. S5b).

Assuming that a single swarm trajectory drifts in CV space with a vector, $\mathbf{s}$, from its starting point on the string, the average drift vector, $\mathbf{S}$, for $n$ swarm trajectories is:

$$\mathbf{S}(n) = \frac{\sum_{i=1}^{n} \mathbf{s}_i}{n} \tag{1}$$

We suggest a convergence metric, $\Delta S$, to estimate how well $\mathbf{S}$ has converged:

$$\Delta S(n) = \frac{|\mathbf{S}(n) - \mathbf{S}(n-b)|}{|\mathbf{S}(n)|} \tag{2}$$

where $b \geq 1$ is the *batch size*, equal to the number of simulations running in parallel on a single compute node. As long as $\Delta S(n)$ is above a certain threshold, batches of additional $b$

short trajectories are launched from that point (Fig. S4). The iteration stops when $\Delta S(n)$ reaches the chosen threshold.

##### Optimized string reparametrization between iterations

The points along the string do not necessarily need to lie equidistant in CV space; what is important is to prevent all the points from drifting to the nearest free energy minima and instead force some points to lie in transition regions of high free energy. The re-positioning of the points along the string is broadly called reparametrization (Fig. S4). In order to make sure that the we sample in all regions along the string, we propose to the keep the points at a distance such that the fraction of swarm trajectories transitioning from one point to a neighbouring point is approximately constant. To do so, relative weights proportional to the length of the arc between two adjacent points on the string are added to the reparametrization algorithm. The weights are estimated from the drift of the swarms according to the following scheme for two points $i$ and $j = i + 1$:

1. For every swarm trajectory, $\zeta$, starting from point $i$, compute the distance in CV space from the final coordinates of the trajectory $\mathbf{x}_\zeta$ to the coordinates of the points $\mathbf{x}_i$ and $\mathbf{x}_j$ where $j = i + 1$. Define the transition metric $\gamma_{i\to j,\zeta} = \frac{|\mathbf{x}_\zeta - \mathbf{x}_i|}{|\mathbf{x}_\zeta - \mathbf{x}_i| + |\mathbf{x}_\zeta - \mathbf{x}_j|}$. $\gamma_{i\to j,\zeta} = 1$ means a full transition from $\mathbf{x}_i$ to $\mathbf{x}_j$ and 0 means no transition at all. Also compute $\gamma_{j\to i,\zeta}$.

2. Compute the average forward and backward transition metrics: $\gamma_{i\to j} = \frac{\sum_\zeta \gamma_{i\to j,\zeta}}{\sum_\zeta}$ and $\gamma_{i\leftarrow j} = \frac{\sum_\zeta \gamma_{j\to i,\zeta}}{\sum_\zeta}$.

3. Compute the transition weights, $w_i$, one per arc on the string: $w_i = \max(\epsilon, \gamma_{i\leftarrow i+1}, \gamma_{i\to i+1})$. $\epsilon$ is a small number to avoid zero weights, which would correspond to two points having the exact same coordinates.

4. Normalize the elements of the weight vector $\mathbf{w}$ so that $\sum_i w_i = 1$.

5. Finally, reparametrize the string so that the distance of the arc between point $i$ and point $i+1$ is $w_i L$, where $L$ is the total length of the string. We see that setting all weights to a constant value corresponds to keeping all arcs equidistant.

Another modification to the algorithm is to gradually increase the string resolution along the minimum path optimization. This will increase the sampling intensity in the vicinity of the string around the most probable transition path, eventually yielding a higher resolution of the free energy landscape. The number of points on the string is changed at defined iterations by inserting an extra point at the start of the string and reparametrizing it again.

##### Configuration exchange between string points enables smoothing orthogonal CVs

The most straightforward way to set up a new iteration of the string simulation is to copy the output of the restrained simulation for a certain point and use as initial coordinates for the next iteration's restrained simulation for the same point. However, this may result in unrestrained orthogonal degrees of freedom getting trapped in local free energy minimas. In theory, two adjacent points may end up structurally quite different in all unbiased dimensions. We therefore use the closest output coordinates in CV space of any trajectory from the previous iteration as input for the restrained simulation (Fig. S5c). This increases sampling of the orthogonal degrees of freedom and also allows us to run the restrained simulations for a shorter period of time, since the input coordinates are already optimally close to the center of the harmonic potential. The optimized string-of swarms simulation protocol was implemented in python. The MDTraj python library (*37*) was used to load and analyze trajectories.

#### Free Energy Computation

The string does not converge to a unique configuration; it diffuses between an equilibrium of minimum free energy pathways and the swarms can be considered to come from a stochastic diffusion process at equilibrium. By discretizing the system into microstates, or bins, it is possible to use the short trajectories from the swarms to create a transition matrix and derive the free energy distribution of the system (*38*) along some variable (Fig. S9).

The transition probabilities $T_{ij}$ of the transition matrix $T$ can be estimated from the normalized number of transitions, $N_{ij}$, from bin $i$ to bin $j$:

$$T_{ij} = \frac{N_{ij}}{\sum_k N_{ik}} \tag{3}$$

The transition matrix of a physical system at equilibrium is constrained by detailed balance, such that for the stationary probability distribution, $\rho$:

$$\rho_i T_{ij} = \rho_j T_{ji} \tag{4}$$

However, due to the limited amount of sampling from the swarms, the resulting transition matrix does not necessarily fulfill this condition. Instead, we have to regularize the transition matrix to obey detailed balance in the same way as is typically done to construct a reversible Markov State Model (*39*). The eigenvector with unit eigenvalue of such a matrix is the stationary distribution, $\rho$.

The free energy, $\Delta G$ of the system is simply computed with the Boltzmann factor:

$$\Delta G_i = -k_B T \log \rho_i \tag{5}$$

In this work, the swarm trajectories from the last 200 iterations were used to construct the transition matrix and to compute the free energy along the variables described in Table S2.

**Even though Eq. 4 is valid for any variable and timescale, the dynamics of the system can be affected by projection onto a lower-dimensional space and may be sensitive to short lagtimes (i.e. the length of the swarm trajectories in this context) when constructing the transition matrix. A common test of convergence is to compute the timescales for a sequence of lagtimes and select the shortest**

**lagtime for which the kinetics do not change(*39*). Such test is not possible to perform with the current implementation due to the use of short single-frame swarm trajectories. See Ref. *40* for a discussion on how the results of the string method can be used as a first step to construct Markov state models.**

One should bear in mind that the resulting free energy landscape can be slightly affected by a the choice of discretization technique. In this paper figures are generated with bins of fixed width. If the bins are too large it will lead to free energy barriers of reduced height. The reason for this is that the barrier may not be resolved properly and mixed with neighbouring regions of lower free energy. On the other hand, a bin width too small will lead to unphysical oscillations in the free energy profile due to insufficient sampling.

##### Error estimation from a posterior distribution of transition matrices

The number of transitions is finite and varies with the size of the system. Therefore there is an uncertainty associated with each transition probability $T_{ij}$. Since many elements go into constructing $T$, the impact of all the statistical uncertainties on the final free energy landscape cannot be directly derived. Instead, we apply a Bayesian approach and use Metropolis Markov chain Monte Carlo (MCMC) to sample over the posterior distribution of transition matrices, given the elements of $T_{ij}$ (*39*). As a result we obtain a number of transition matrices and can directly compute the mean and variance of the stationary probability distribution, and therefore also of the free energy landscape. Thus, this gives us another way to assert convergence since trajectories drawn from an equilibrium ensemble will generate a more narrow distribution of free energy profiles than trajectories drawn from different non-equilibrium processes. We estimated the errors on the free energies with a MCMC method using the implementation of a regularized transition matrix in MSMBuilder (*39*). In all error estimations, we sampled 1000 transition matrices.

Outliers in the distribution of sampled free energy profiles (Fig. S8, S10 and S11) may in a few cases reverse the relative stability of states. In general, however, the majority of the the free energy values within the upper and lower whiskers all give a similar view of the relative stability between states for different conditions.

13. Che, T., Majumdar, S., Zaidi, S. A., Ondachi, P., McCorvy, J. D., Wang, S., Mosier, P. D., Uprety, R., Vardy, E., Krumm, B. E., Han, G. W., Lee, M.-Y., Pardon, E., Steyaert, J., Huang, X.-P., Strachan, R. T., Tribo, A. R., Pasternak, G. W., Carroll, F. I., Stevens, R. C., Cherezov, V., Katritch, V., Wacker, D., and Roth, B. L.

(2018) Structure of the Nanobody-Stabilized Active State of the Kappa Opioid Receptor. *Cell 172*, 55–67.e15.

20. Yin, W., Zhou, X. E., Yang, D., de Waal, P. W., Wang, M., Dai, A., Cai, X., Huang, C.-Y., Liu, P., Wang, X., Yin, Y., Liu, B., Zhou, Y., Wang, J., Liu, H., Caffrey, M., Melcher, K., Xu, Y., Wang, M.-W., Xu, H. E., and Jiang, Y. (2018) Crystal structure of the human 5-HT1B serotonin receptor bound to an inverse agonist. *Cell Discovery 4*.

26. Huang, J., Rauscher, S., Nawrocki, G., Ran, T., Feig, M., de Groot, B. L.,

Grubmüller, H., and MacKerell Jr, A. D. (2016) CHARMM36m: an improved force field for folded and intrinsically disordered proteins. *Nat. Methods 14*, 71–73.

27. Jorgensen, W. L., Chandrasekhar, J., Madura, J. D., Impey, R. W., and Klein, M. L. (1983) Comparison of simple potential functions for simulating liquid water. *J. Chem. Phys. 79*, 926–935.

28. Kim, S., Lee, J., Jo, S., Brooks III, C. L., Lee, H. S., and Im, W. (2017) CHARMM-GUI ligand reader and modeler for CHARMM force field generation of small molecules. *J. Comput. Chem. 38*, 1879–1886.

29. Abraham, M. J., Murtola, T., Schulz, R., Páll, S., Smith, J. C., Hess, B., and Lindahl, E. (2015) GROMACS: High performance molecular simulations through multi-level parallelism from laptops to supercomputers. *SoftwareX 1*, 19–25.

30. Tribello, G. A., Bonomi, M., Branduardi, D., Camilloni, C., and Bussi, G. (2014) PLUMED 2: New feathers for an old bird. *Comput. Phys. Commun. 185*, 604–613.

31. Schrödinger, LLC, (2015) The PyMOL Molecular Graphics System, Version 2.1.

32. Dror, R. O., Arlow, D. H., Maragakis, P., Mildorf, T. J., Pan, A. C., Xu, H., Borhani, D. W., and Shaw, D. E. (2011) Activation mechanism of the $\beta$2-adrenergic receptor. *Proc. Natl. Acad. Sci. U.S.A. 108*, 18684–18689.

33. Ponzoni, L., Polles, G., Carnevale, V., and Micheletti, C. (2015) SPECTRUS: A dimensionality reduction approach for identifying dynamical domains in protein complexes from limited structural datasets. *Structure 23*, 1516–1525.

34. Westerlund, A. M., and Delemotte, L. (2018) Effect of Ca2+ on the promiscuous target-protein binding of calmodulin. *PLOS Comput. Biol. 14*, 1–27.

35. Pedregosa, F., Varoquaux, G., Gramfort, A., Michel, V., Thirion, B., Grisel, O., Blondel, M., Prettenhofer, P., Weiss, R., Dubourg, V., Vanderplas, J., Passos, A., Cour-

napeau, D., Brucher, M., Perrot, M., and Duchesnay, E. (2011) Scikit-learn: Machine Learning in Python. *J. Mach. Learn. Res. 12*, 2825–2830.

36. Montavon, G., Lapuschkin, S., Binder, A., Samek, W., and Müller, K.-R. (2017) Explaining nonlinear classification decisions with deep taylor decomposition. *Pattern Recognit. 65*, 211–222.

37. McGibbon, R. T., Beauchamp, K. A., Harrigan, M. P., Klein, C., Swails, J. M., Hernández, C. X., Schwantes, C. R., Wang, L.-P., Lane, T. J., and Pande, V. S. (2015) MDTraj: A Modern Open Library for the Analysis of Molecular Dynamics Trajectories. *Biophys. J. 109*, 1528 – 1532.

38. Lev, B., Murail, S., Poitevin, F., Cromer, B. A., Baaden, M., Delarue, M., and Allen, T. W. (2017) String method solution of the gating pathways for a pentameric ligand-gated ion channel. *Proc. Natl. Acad. Sci. U.S.A. 114*, E4158–E4167.

### Supplementary Figures

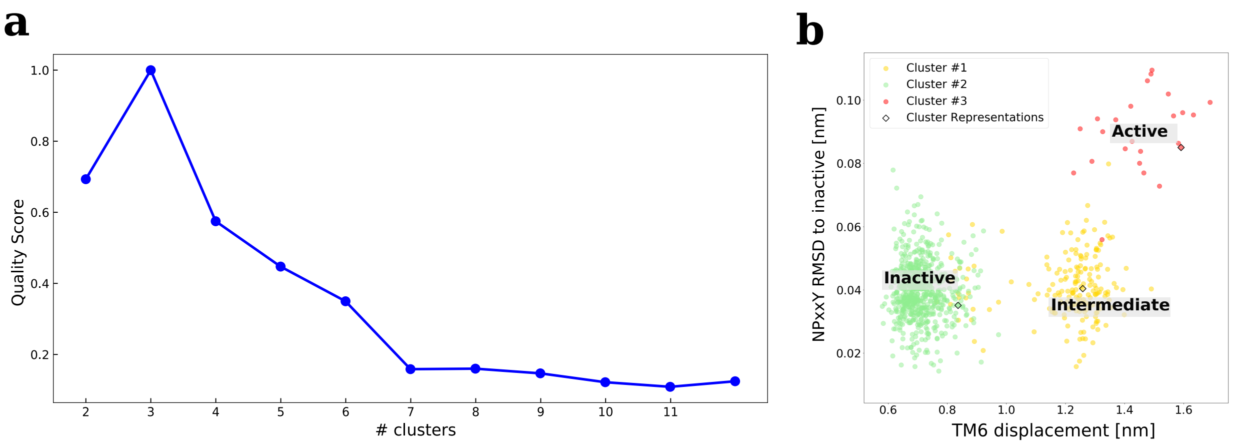


Figure S1: Clustering of a long MD simulation trajectory of $\beta_2$AR deactivation into states (*32*). (**a**) The SPECTRUS quality score for different number of clusters, normalized such that the highest value is 1. The peak occurs at 3 clusters, indicating the data is optimally partitioned into three clusters. (**b**) Three states found via spectral clustering on the C$\alpha$ inter-residue inverse distance map. Each configuration is colored according to its cluster assignment and projected in the TM6 displacement and NPxxY RMSD to inactive space. The cluster centers (configurations with minimum total RMSD to all other configuration in that cluster) are marked as empty circles.

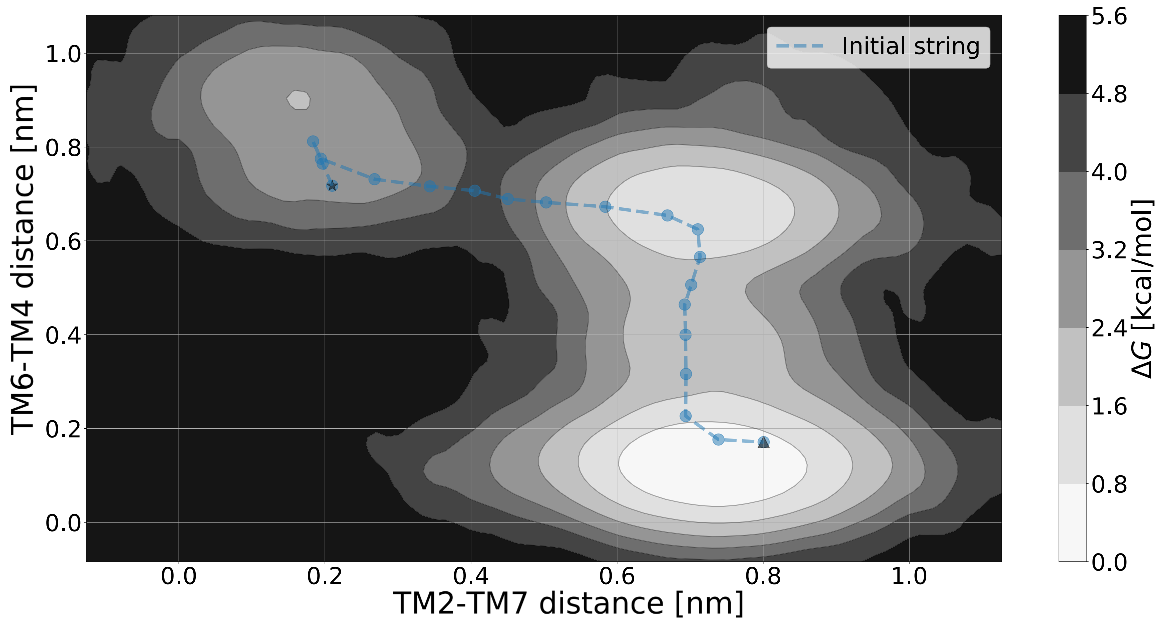


Figure S2: Initial path guess, via rough estimation of the free energy landscape corresponding to configurations from a long time scale trajectory (*32*). The density map that serves as a basis to compute the free energy landscape is computed by a kernel density estimator (KDE) and projected onto the top two original CVs. The fixed starting and end points, marked with a star and a triangle respectively, are taken to be the equilibrated crystal structures 3P0G and 2RH1.

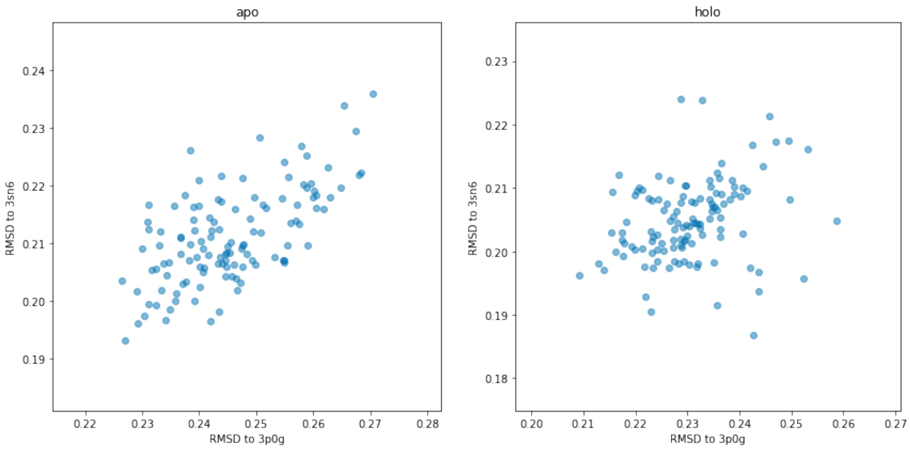


Figure S3: Root-mean-square-deviation (RMSD) to nanobody bound active structure 3P0G (x-axis) and to the G protein bound structure 3SN6 (y-axis). The points correspond to the restrained output coordinates for every point the active substring from the last 10 iterations. (**a**) for the apo receptor and (**b**) for the holo receptor.

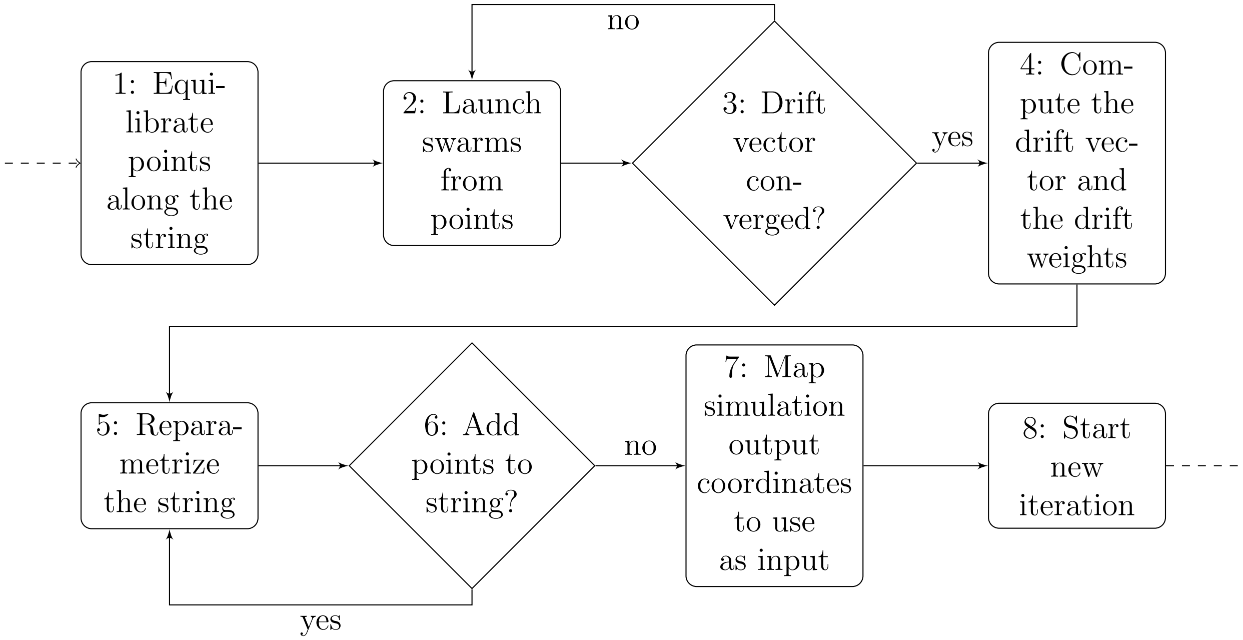


Figure S4: An optimized iteration along the string-of-swarms algorithm. The input frames from the previous iteration are equilibrated under CV-based restraints (box 1). The swarm trajectories are then launched in parallel batches from the points (box 2). As the short trajectories finish, the drift vector is computed and updated. If the vector changes above a threshold (diamond 3), new batches will be launched (Fig. S5b). Once the drift vector has converged, the final drift as well as the drift weights, a measure of how much the swarm trajectories tend to move between two neighboring points, are computed (box 4). The points are then redistributed (so-called reparametrized) on the string (box 5), such that the number of transitions between neighboring points is kept relatively constant. At selected iterations, new points are also added to the string (diamond 6), requiring additional reparametrization (Fig. S5a). The configurations with coordinates closest to the new points of the reparametrized string are used as input for the next iteration (box 8) (Fig. S5c).

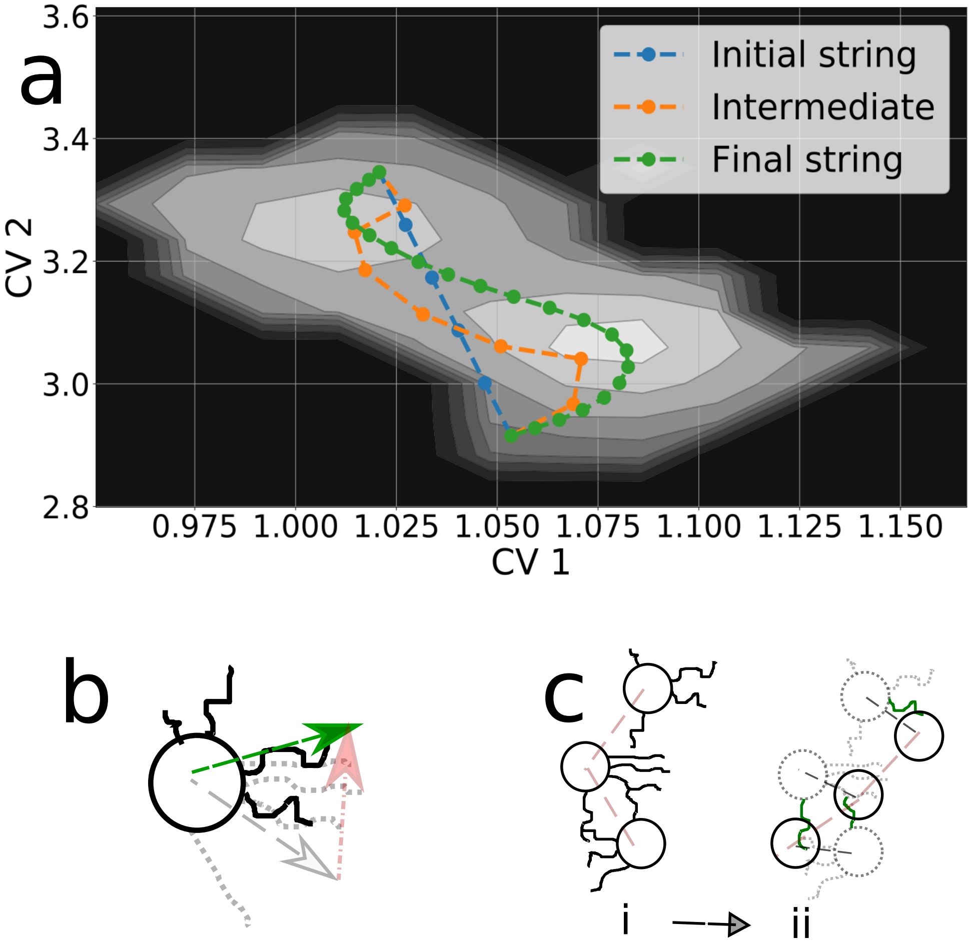


Figure S5: Algorithmic optimization of the string method with swarms of trajectories. (**a**) The method allows to iteratively find the most probable transition path between two fixed endpoints, as illustrated here for a simple 2 well model. To increase the resolution of the minimum free energy path as a minimal computational cost, the number of points on the string, and from which the swarms are launched, is gradually incremented and the lengths of the arcs on the strings are set to maximize the transitions between neighbouring points. (**b**) The number of swarms per string point is optimized on-the-fly by designing an adaptive scheme: the swarm convergence is evaluated by computing the difference between the average drift vector before (grey) and after (green) the last batch of short trajectories (black trajectories) has finished; if the norm of the difference (red) divided by the norm of the average drift after is above a certain limit, new trajectories are instantiated. This is done until this difference reaches a value before the defined limit. (**c**) To promote convergence and prevent different string points from diverging from one another in the dimensions that were not considered in the CV space, exchanges between string points are promoted: to select input coordinates for the next string iteration (black circles in **ii**), configurations from the closest swarm trajectory (green trajectories in **ii**) originating from the previous string iteration (black circles in **i**, grey circles in **ii**) are selected. These can come from the same string point or from a neighboring one.

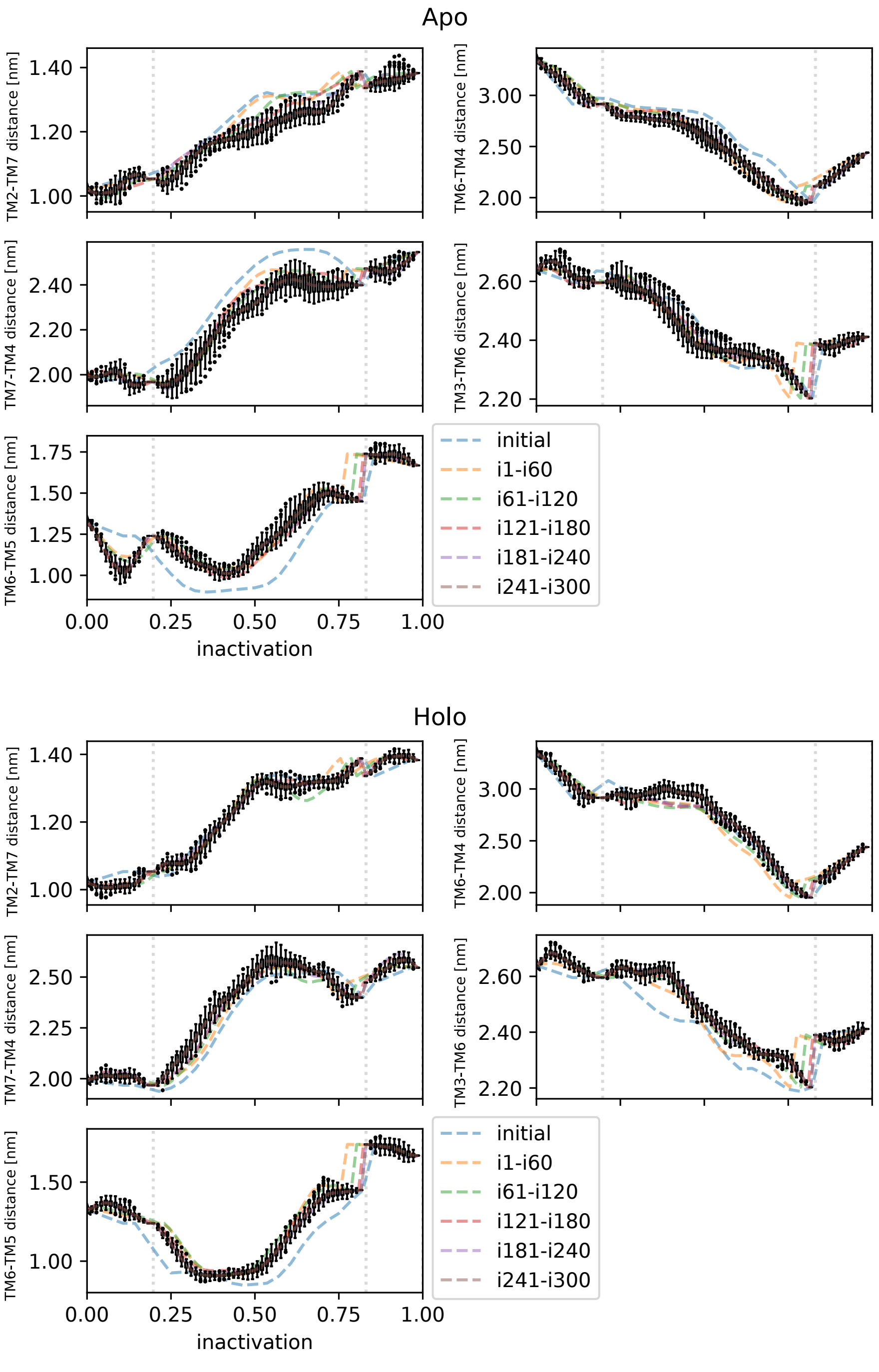


Figure S6: Average strings for different iterations, projected along the original CVs, for conditions Apo and Holo. The x axis shows progression along the string from the active towards the inactive state. The whiskers show the interquartile range for the final 60 iterations, outliers are shown as circles. The active, transition and inactive substrings are separated by a dashed vertical line.

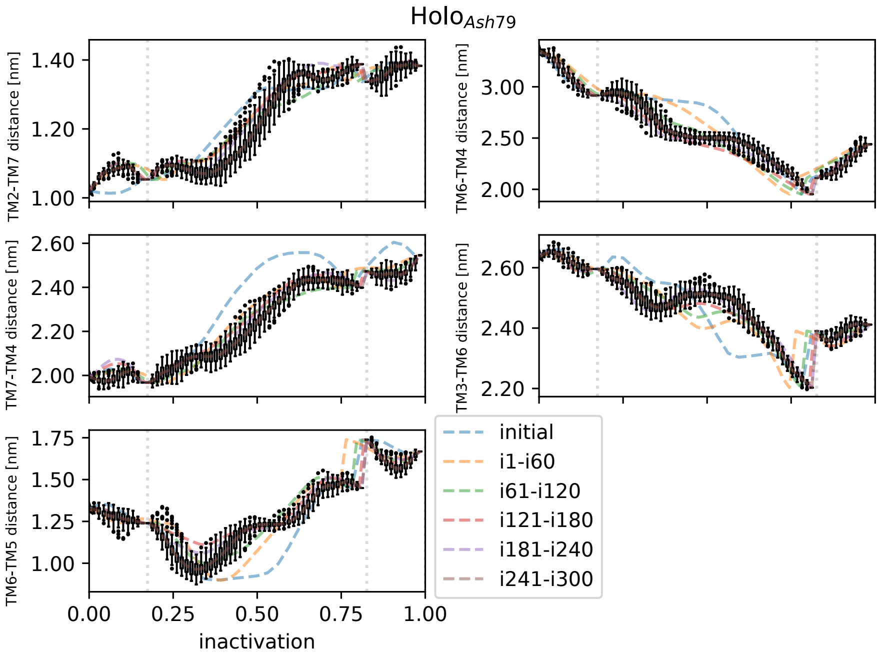


Figure S7: Average strings for different iterations, projected along the original CVs, for condition Holo$_{Ash79}$. The x axis shows progression along the string from the active towards the inactive state. The whiskers show the interquartile range for the final 60 iterations, outliers are shown as circles. The active, transition and inactive substrings are separated by a dashed vertical line.

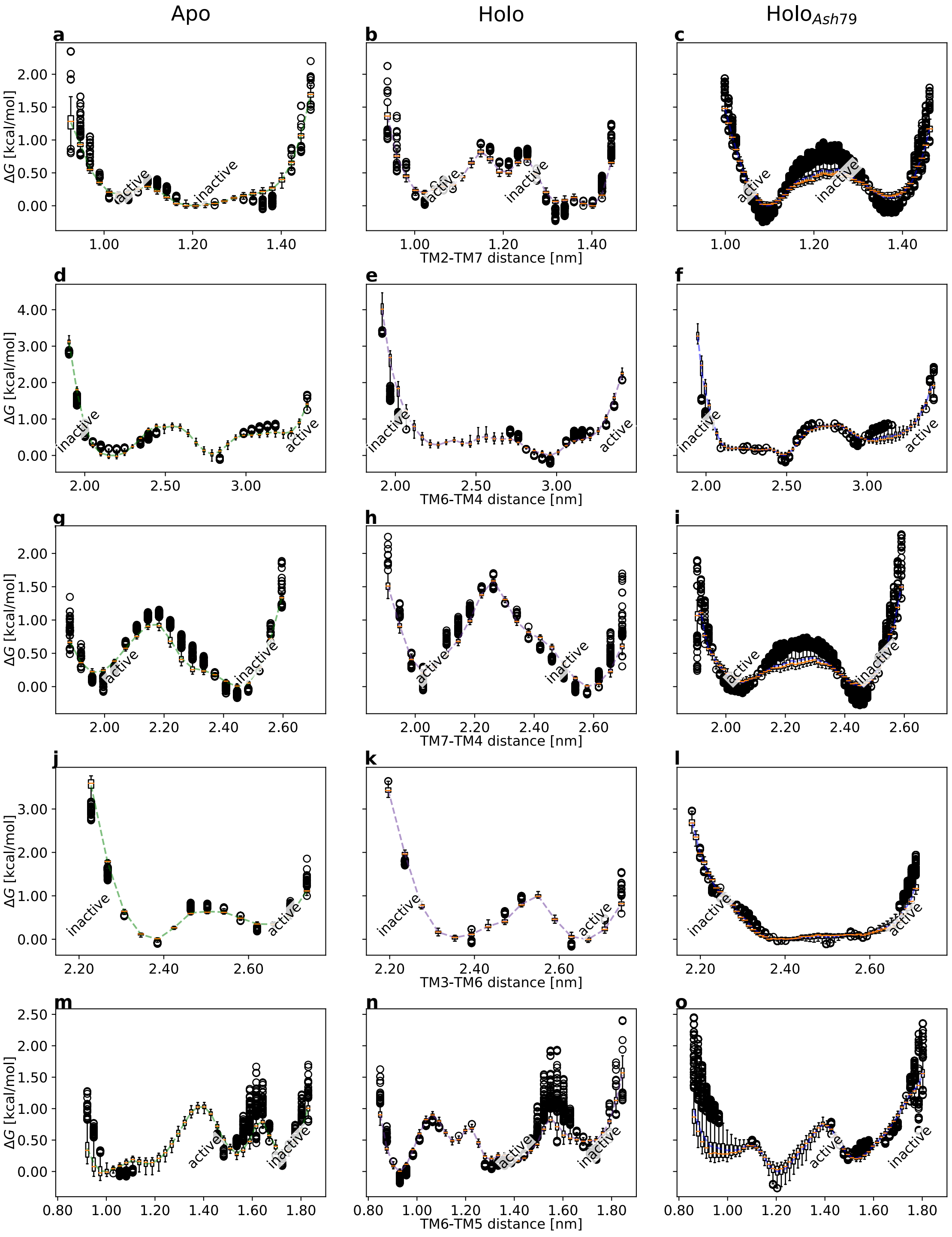


Figure S8: Distribution of free energy landscapes from a MCMC sampling round over 1000 transition matrices projected onto the original CVs. The boxplots show the median (as a horizontal line in orange), the interquartile range (as a box), the upper and lower whiskers (as vertical lines) as well as the outliers (as circles).

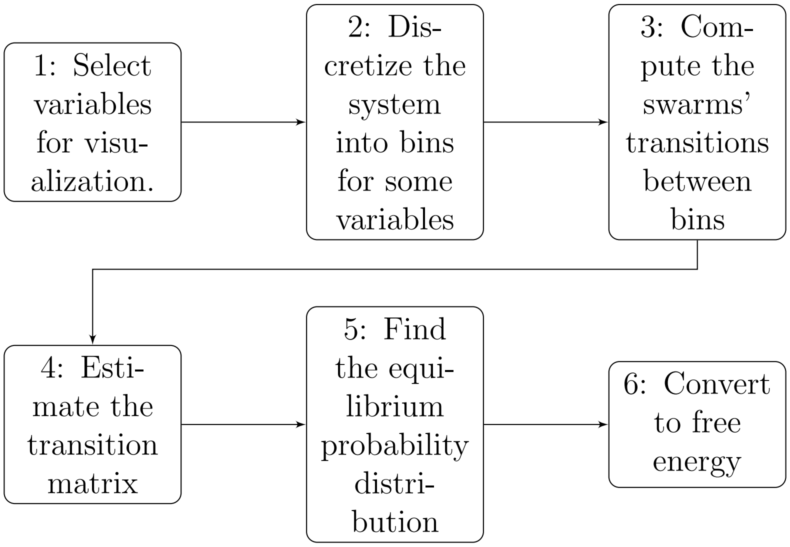


Figure S9: Postprocessing protocol for computing the free energy in the vicinity of the activation path. Given some variable of interest to project the free energy onto (box 1), the system is partitioned into bins along this variable (box 2). Next, we iterate through all swarm trajectories and count the number of transitions between all bins (box 3). A transition probability matrix is then constructed (box 4). We then compute the probability distribution at equilibrium (box 5), i.e. the probability for the system to be in a certain bin while fulfilling detailed balance (Eq. 4). Finally, the free energy of all bins is computed from the probability distribution.

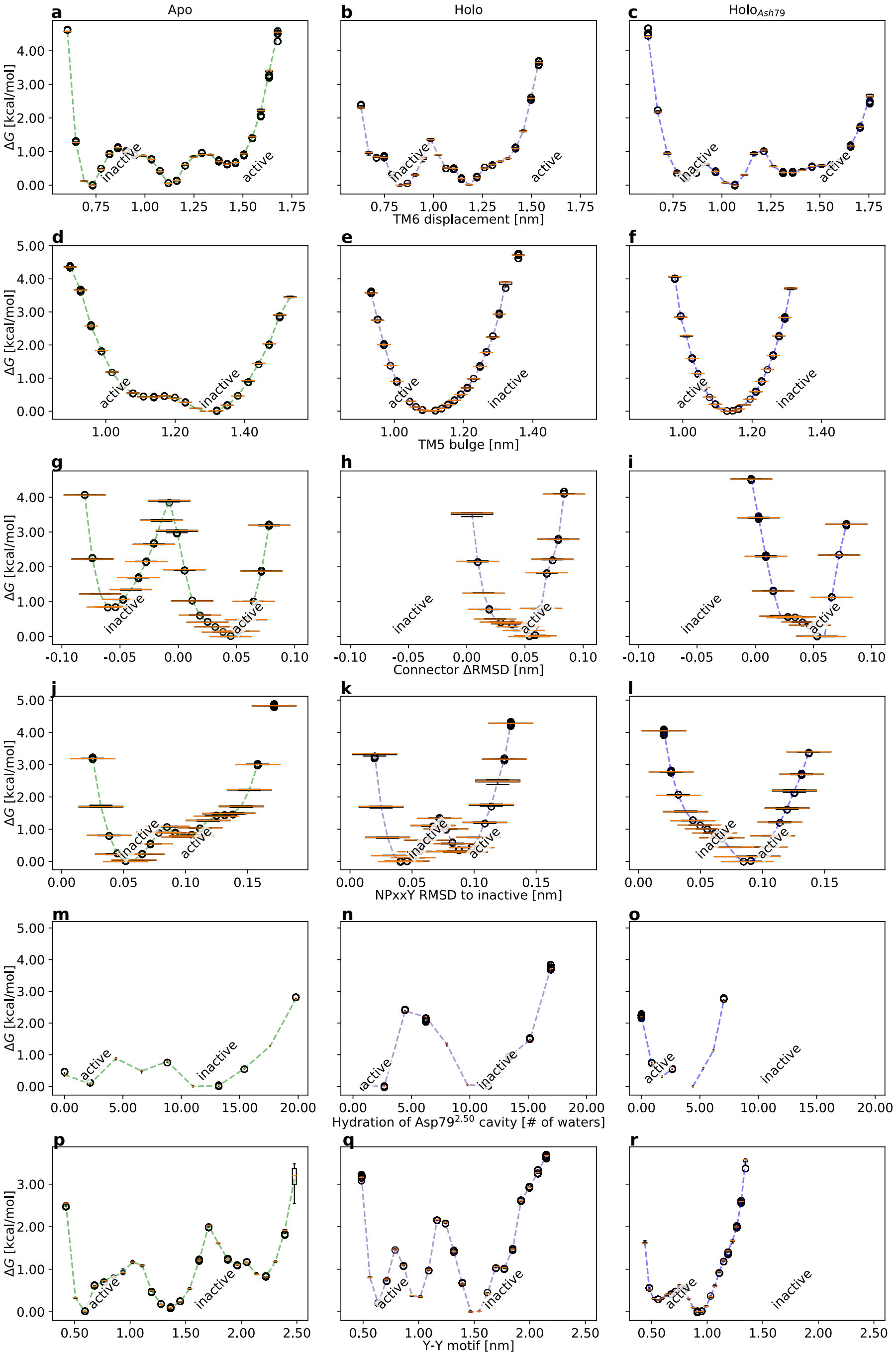


Figure S10: Uncertainties associated with the free energy landscapes projected onto various variables of interest (displacement of TM6, distance between Ser120$^{3.39}$ and Gly315$^{7.41}$, connector change in RMSD, twist of the NPxxY motif, hydration of the Asp79$^{2.50}$ cavity, the YY-motif) from an MCMC sampling round over 1000 transition matrices. The boxplots show the median (as a horizontal line in orange), the interquartile range (as a box), the upper and lower whiskers (as vertical lines) as well as the outliers (as circles).

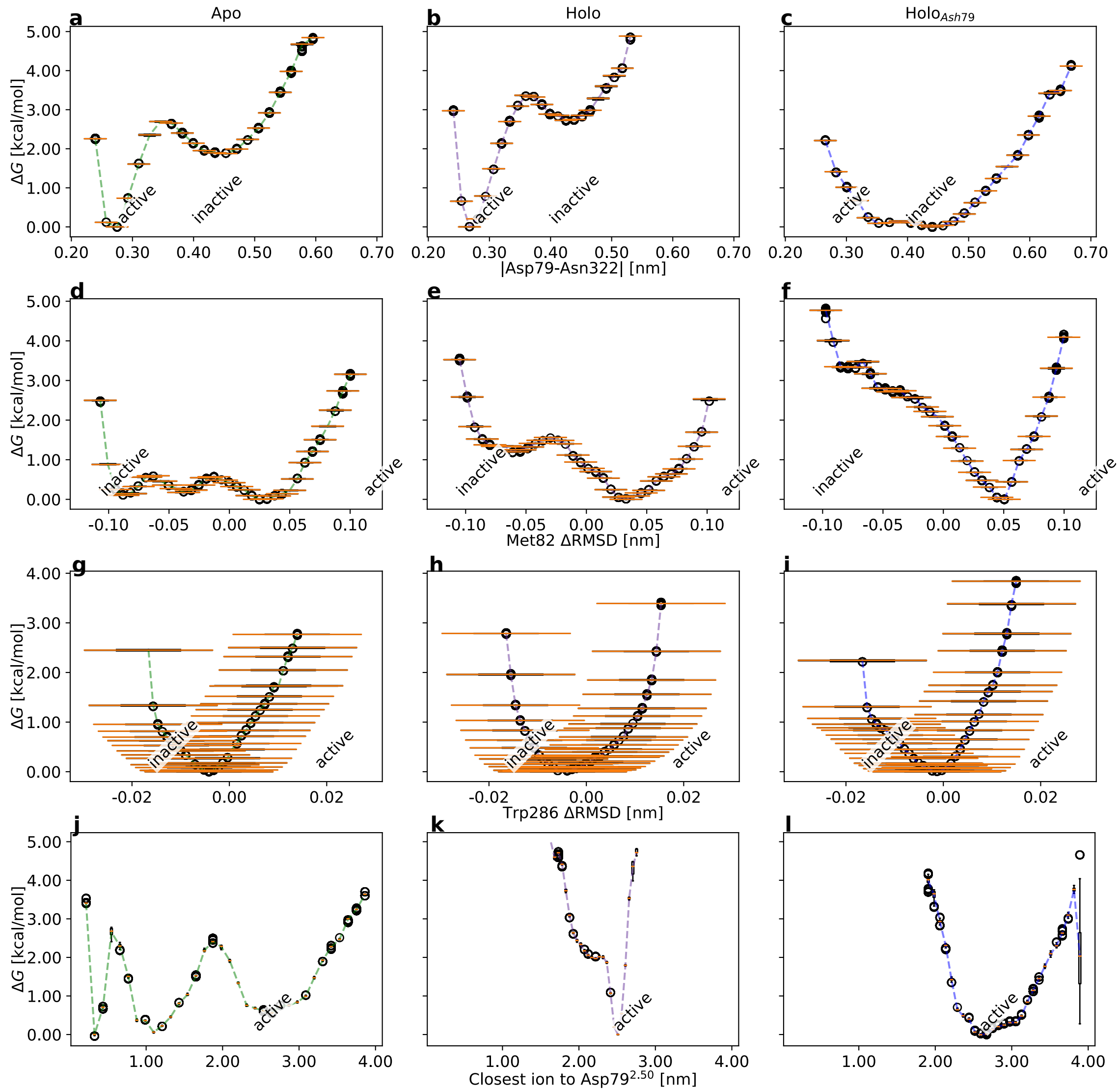


Figure S11: Uncertainties associated with the free energy landscapes projected onto various variables of interest including the distance between Asp79$^{2.50}$ and Asn322$^{7.49}$ (**a-c**), the Met82$^{2.53}$ heavy atom ΔRMSD) (**d-f**), the Trp286$^{6.48}$ (the so-called 'toggle-switch' residue) heavy atom ΔRMSD) (**g-i**) **and the distance from Asp79$^{2.50}$ to the closest sodium ion (j-l)** from an MCMC sampling round over 1000 transition matrices. The boxplots show the median (as a horizontal line in orange), the interquartile range (as a box), the upper and lower whiskers (as vertical lines) as well as the outliers (as circles).

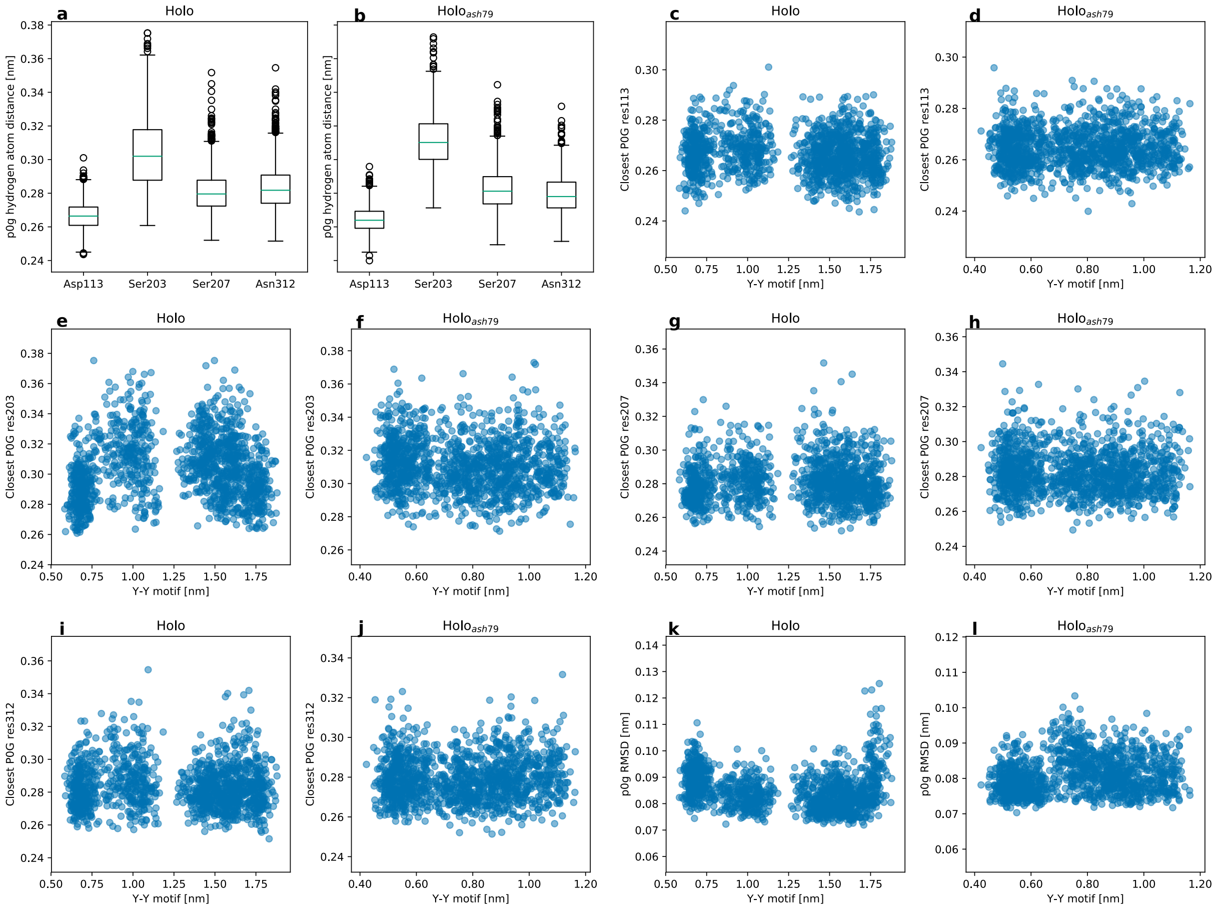


Figure S12: (**a-b**) Boxplots of the closest heavy atom distance from Asp113$^{3.32}$, Ser203$^{5.43}$, Ser207$^{5.46}$ and Asn312$^{7.38}$ to the ligand BI-167107 (P0G). The whiskers show the interquartile range and outliers are shown as circles. (**c-l**) Y-Y motif (x-axis) plotted against different variables of the binding site: closest interaction distance from the ligand to Asp113$^{3.32}$ (**c-d**), Ser203$^{5.43}$ (**e-f**), Ser207$^{5.46}$ (**g-h**) and Asn312$^{7.38}$ (**i-j**) respectively. (**k-l**) Root-mean-square-deviation (RMSD) to the ligand from the active structure 3P0G (y-axis). The output coordinates from the swarm and restrained trajectories of the final iteration have been used as datapoints in all figures. For the ligand bound simulations with Asp79$^{2.50}$ ionized (**a**, **c**, **e**, **g**, **i** and **k**) or protonated (**b**, **d**, **f**, **h**, **j** and **l**).

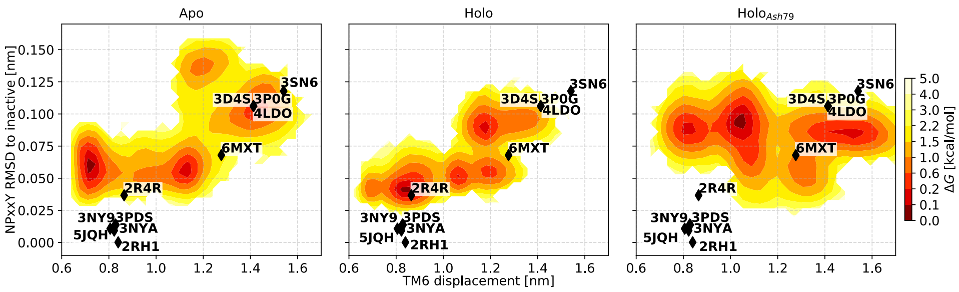


Figure S13: Free energy of the TM6 displacement and the NPxxY RMSD to the inactive structure 2RH1. The coordinates of $\beta_2$AR structures (3P0g(*23*), 3SN6(*1*), 4LDO(*41*), 3D4S(*42*), 6MXT(*43*), 2R4R(*44*), 5JQH(*45*), 3NYA(*46*), 3NY9(*46*), 3PDS(*47*) and 2RH1 (*2*)) are shown with their corresponding PDB IDs.
